# Sequential addition of neuronal stem cell temporal cohorts generates a feed-forward circuit in the Drosophila larval nerve cord

**DOI:** 10.1101/2022.04.05.487221

**Authors:** Yi-wen Wang, Chris C Wreden, Maayan Levy, Zarion D Marshall, Jason N MacLean, Ellie S Heckscher

## Abstract

Understanding how circuits self-assemble starting from neuronal stem cells is a fundamental question in developmental biology. Here, we addressed how neurons from different lineages wire with each other to form a specific circuit motif. To do so, we combined developmental genetics—Twin spot MARCM, Multi-color Flip Out, permanent labeling—with circuit analysis—calcium imaging, connectomics, and network science analyses. We find many lineages are organized into temporal cohorts, which are sets of lineage-related neurons born within a tight time window, and that temporal cohort boundaries have sharp transitions in patterns of input connectivity. We identify a feed-forward circuit motif that encodes the onset of vibration stimuli. This feed-forward circuit motif is assembled by preferential connectivity between temporal cohorts from different neuronal stem cell lineages. Further, connectivity does not follow the often-cited early-to-early, late-to-late model. Instead, the feed-forward motif is formed by sequential addition of temporal cohorts, with circuit output neurons born before circuit input neurons. Further, we generate multiple new tools for the fly community. Ultimately, our data suggest that sequential addition of neurons (with outputs neurons being oldest and input neurons being youngest) could be a fundamental strategy for assembling feed-forward circuits.

## Introduction

Neuronal circuits are the fundamental functional units of the nervous system, and neuronal stem cell lineages are fundamental developmental units. Determining lineage-circuit relationships is essential for deciphering the developmental logic of circuit assembly (Li et al., 2018)(Meng & Heckscher, 2021). Circuit membership of lineage-related neurons have so far been mapped in an ad-hoc, lineage-by-lineage basis in a handful of model circuits. For example, in the vertebrate neocortex, excitatory neurons from a single stem cell preferentially populate individual neocortical microcolumns (Y.-C. Yu et al., 2009). In the vertebrate hippocampus, excitatory neurons from a single stem cell share inhibitory input (Xu et al., 2014). In the Drosophila visual system, synchronous production of neurons from a single stem cell underlie the retinotopy of direction-selective neurons (Pinto-Teixeira et al., 2018). Together these studies demonstrate that lineage-circuit relationships differ depending on brain region and stem cell type examined. However, it remains unknown how neurons from different stem cell lineages wire with each other to form specific circuit motifs. In this study, we took a new approach, the converse of previous studies. Here we start from circuit motif and identify the developmental origins of neurons of the circuit. This approach allowed us to begin to identify the developmental rules that govern the self-assembly of circuits starting from stem cells.

As a model, we use the Drosophila larval nerve cord. Nerve cord circuits, like those in spinal cord, processes multiple somatosensory stimuli and generate patterned muscle contractions (Meng & Heckscher, 2021). Spinal/nerve cords are bilaterally symmetric and segmented. They contain motor neuron and interneuron cell types that are homologous to each other (Heckscher et al., 2015)(Catela & Kratsios, 2021). However, only for the Drosophila larval nerve cord is a connectome available. This connectome is a high-resolution, transmission electron microscopic image volume of all neurons and synapses in the CNS (Ohyama et al., 2015b). It enables anatomical circuit tracing at cellular and synaptic resolution. Furthermore, for the Drosophila nerve cord, neuronal stem cells (neuroblasts), the lineages of neurons they generate and the molecular control of neuronal diversity are among the best-characterized in biology (C. Q. Doe, 2017). In each Drosophila larval nerve cord segment, spatial patterning transcription factors (e.g., row and column genes) establishes 30 unique neuroblast types (*Figure 1A*) (J. J. Broadus et al., 1995). Each neuroblast produces a unique, invariant lineage of neurons (e.g., *Figure 1B)(Schmid et al., 1999)*. To do so, the neuroblast divides to self-renew and to generate a ganglion mother cell (GMC). The GMC then divides to produce two neurons (*Figure 1B)*, which are differentiated from each other by Notch signaling (J. B. S. and C. Q. Doe, 1998). This process repeats and all Notch ON neurons from a single neuroblast populate an A hemilineage, and all Notch OFF neurons populate a B hemlineage (Truman et al., 2010). This mode of neuroblast division is referred to as a !type 1”. However, some neuroblasts can switch to a “type 0” division mode (e.g., *Figure 1B)* in which the neuroblast directly produces only Notch OFF neurons (Baumgardt et al., 2014). Within a hemilineage, each neuron has a unique temporal identity, or birth-order (e.g., first-born, second-born, etc.), which is specified by temporal transcription factor expression in neuroblasts (Isshiki et al., 2001). It has been suggested that these genetic mechanisms specifying neuronal diversity are also likely to govern circuit formation and function (Mark et al., 2021)(Sagner & Briscoe, 2019).

**Figure 1.**
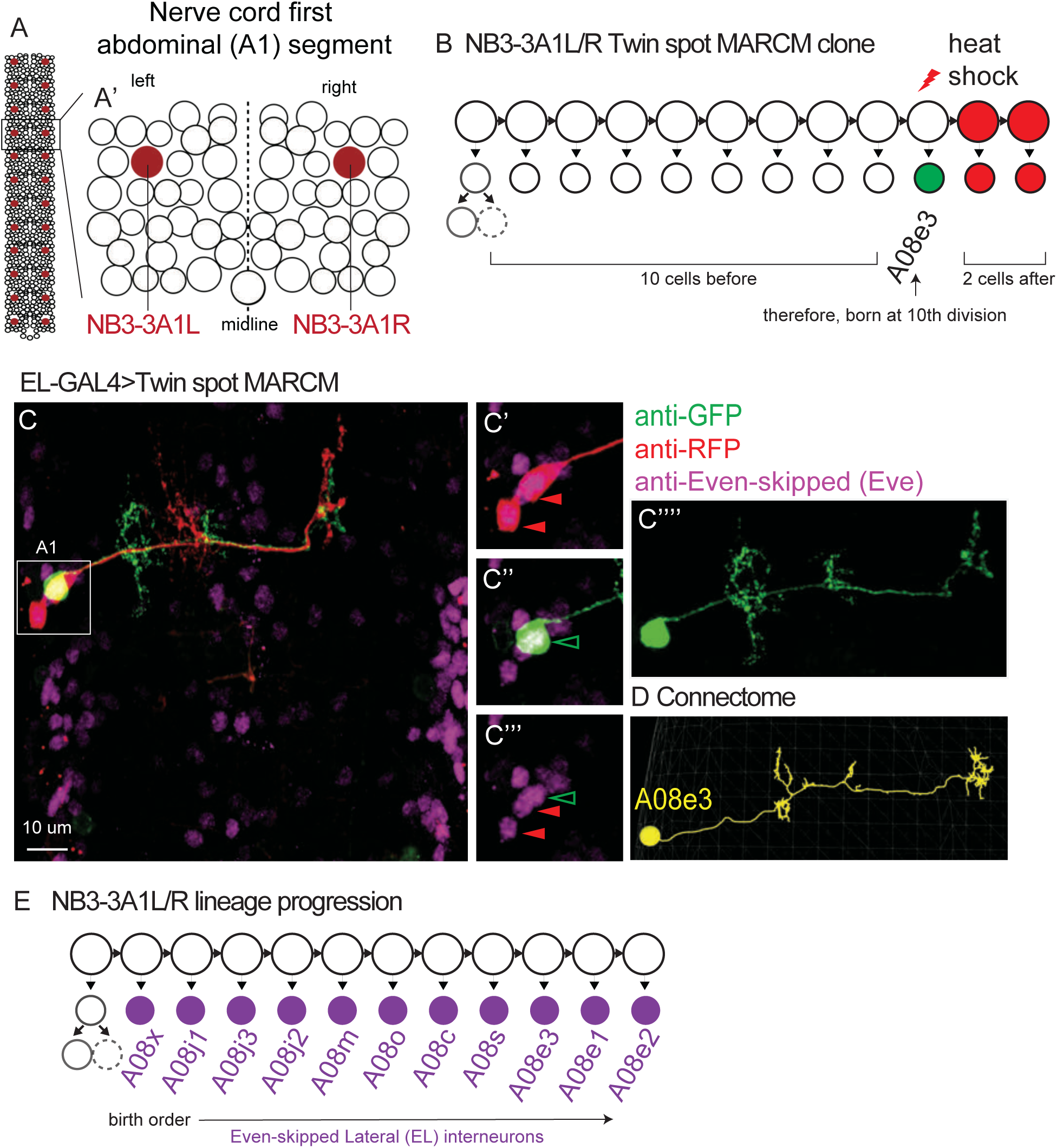
ts-MARCM determines the birth order and morphology of all NB3-3A1L/R neuronal progeny. **A-B. Illustrations of Drosophila neuroblasts** A. The nerve cord is left-right symmetrical and segmented. Each circle represents one neuroblast with NB3-3 in maroon. Segment A1 (boxed) is enlarged in A’. It contains 30 types of neuroblasts. B. NB3-3 lineage progression is shown with an example ts-MARCM clone overlaid. Each circle represents one cell and each arrow represents a cell division. First, NB3-3 divides to self-renew and generate a ganglion mother cell, which divides to generate a motor neuron (solid circle) and an undifferentiated cell (dashed circle). Then, NB3-3 directly generate ELs. In ts-MARCM, a heat shock is provided (red lightning bolt) as NB3-3 divides. In this example, a singly labeled neuron is shown in green (A08e3), and two alternatively-labeled neurons are shown in red. Because the total number of neurons in the lineage are known, counting labeled neurons allows inference of neuronal birth order. The identity of the singly-labeled neuron is determined by matching the labeled neuron to the corresponding neuron in the connectome using morphological criteria (see Methods). **C-D. Image of a ts-MARCM clone and a corresponding neuron in the connectome** C. Many segments of the nerve cord are shown in dorsal view with anterior up. The boxed region is enlarged at the right. In this ts-MARCM clone, two neurons are labeled in red and one in green (arrowheads), and all are Eve(+) ELs. The singly-labeled EL is enlarged to highlight morphological detail. And the corresponding neuron in the connectome is shown in D. Specific genotype is listed in Table S4.

Evidence that supports this idea that the genetic mechanisms specifying neuronal diversity govern circuit assembly comes from the identification of “temporal cohorts” in the Drosophila larval nerve cord. Temporal cohorts are sets of neurons in a hemilineage born within a tight time window. Initially, temporal cohorts were correlated with circuit membership using functional approaches. For example, the NB3-3 stem cell generates a series of EL interneurons (<u>E</u>ven-skipped [eve] neurons with <u>L</u>aterally placed cell bodies) (Wreden et al., 2017). Molecularly, ELs can be subdivided into early-born EL and late-born EL temporal cohorts based on the expression of an enhancer called 11F02 (Wreden et al., 2017). Using calcium imaging, EL interneurons of the early-born temporal cohort responded to vibrational stimuli, whereas EL interneurons of the late-born temporal cohort did not (Wreden et al., 2017). Further, using optogenetics, activation of early-born ELs triggered escape rolling, whereas activation of late-born ELs altered left-right symmetrical crawling (Wreden et al., 2017). These data linked temporal cohorts to differential circuit function. More recently, in other lineages (i.e., NB1-2, NB2-1, NB3-1, NB4-1, NB5-2, NB7-1, NB 7-4), neurons within temporal cohorts were shown to share similarities in their synaptic partnerships (Meng et al., 2019)(Meng et al., 2020)(Mark et al., 2021). Furthermore, the number of neurons in lineage that are segregated into a given temporal cohorts can be altered by manipulating temporal factor expression in neuronal stem cells (Meng et al., 2019)(Meng et al., 2020). Thus, temporal cohorts are developmental units that likely to be specified early during neurogenesis in a lineage intrinsic manner by the molecular programs known to generate neural diversity.

Notably, however, previous studies that described temporal cohorts lacked the resolution to distinguish between “graded” and “sharp” wiring transition models. For many neuronal features, such as morphology, gene expression, and neurotransmitter phenotype, there can be sharp changes in lineage progression. For example, a neuroblast can abruptly change from producing motor neurons to interneurons (Meng et al., 2020) or from producing Eve(+) to Eve(−) neurons (Pearson & Doe, 2003). The idea that temporal cohorts have distinct circuit-level function suggests that there are also sharp changes in the patterns of synaptic partnerships formed by neurons in a lineage, and that these sharp changes are correlated with temporal cohort borders. However, available data are also consistent with an alternative, graded transition model. In the graded transition model, during lineage progression, changes in wiring would slowly transition such that neurons with more similar birth times would have more similar synaptic partnerships. Distinguishing between graded and sharp transition models is fundamental for understanding lineage progression in Drosophila neuroblasts lineages.

The overall objective of this paper is to test the hypothesis that Drosophila larval nerve cord circuits are assembled by preferential connectivity between distinct temporal cohorts. Although an attractive hypothesis, there is limited supporting evidence (Meng et al., 2019)(Mark et al., 2021)(Meng et al., 2020). However, before we can address this hypothesis, in the first part of this paper, we needed to distinguish between graded and sharp wiring transition models. Using lineage tracing, connectomics, and network science-based statistical analysis, we find support for a sharp transition model. And so, the first part of this paper brings into alignment the concepts of temporal cohorts, circuit function, and circuit anatomy at single neuron resolution. In the second part of this paper, we use the circuit containing the early-born EL temporal cohort as a model. Using connectomics and calcium imaging, we show that early-born ELs are the output neurons of a feed-forward sensory processing circuit, which encodes the onset of vibrational stimuli. Next, we combine lineage tracing, single neuron labeling approaches, and connectomics to identify the developmental origins for the major interneuron inputs onto early-born ELs. Specifically, for nerve cord interneurons that synapse onto early-born ELs, we identify the stem cell parent, birth order within a lineage, and birth time relative to early-born ELs. Our data support the hypothesis that the feed-forward circuit is assembled by preferential connectivity between distinct temporal cohorts. Further, our data show that circuit outputs (early-born ELs) are born before circuit inputs. Ultimately, these data suggest that sequential addition of temporal cohorts -- with outputs neurons being oldest and input neurons being youngest -- could be a fundamental strategy for assembling feed-forward circuits.

## Results

### NB3-3 lineage contains two temporal cohorts—early-born ELs and late-born ELs, which are distinctive both in morphology and connectivity

The present study aims to understand how neurons from different stem cell lineages wire with each other to form a specific circuit motif. In particular, we wanted to test the hypothesis that Drosophila larval nerve cord circuits are assembled by preferential connectivity among temporal cohorts (Meng & Heckscher, 2021). First, however, we needed to better understand the relationship between temporal cohorts, circuit function, and circuit anatomy. Specifically, we needed to characterize the relationship between neuronal birth order within a lineage and patterns of synaptic connectivity at single neuron resolution. This will allow us to distinguish between graded and sharp wiring transition models (see Introduction).

As a model, we use the NB3-3 lineage in the first abdominal segment (A1) of the nerve cord (*Figure 1A*). In A1, there is a left-right pair of NB3-3 stem cells, which are thought to be identical. We will refer to these neuronal stem cells as NB3-3A1L/R. First, NB3-3A1L/R undergoes a type 1 division, which generates a ganglion mother cell (GMC)(Baumgardt et al., 2014). The GMC divides to produce two neurons of unknown function (*Figure 1B)(*Schmid et al., *1999).* Next, NB3-3 switches to a type 0 division mode and generates EL interneurons directly (*Figure 1B*)(Baumgardt et al., 2014). The birth timing of ELs has been characterized using three different methods (Tsuji et al., 2008)(Wreden et al., 2017)(Mark et al., 2021) nonetheless the exact relationship between neuronal morphology and birth order for neurons in the NB3-3A1L/R lineage remains unknown. Here, we determined the birth order and morphology of every neuron in the NB3-3A1L/R lineage. Using neuron morphologies, we identified matching neurons in the connectome. Using the connectome, we found all upstream synaptic partners. Finally, we analyzed the patterns of wiring using network science approaches.

#### Birth order and morphology of each neuron in the NB3-3A1L/R lineage

First, we needed to determine birth order and morphology of EL neurons at the single neuron level. To do so, we used the gold-standard in the field for lineage tracing, Twin Spot Mosaic Analysis with a Repressible Cell Marker (herein, ts-MARCM) (H.-H. Yu et al., 2009). Briefly, in ts-MARCM heat shocks are delivered to dividing cells, and this renders progeny competent to express either a UAS-red or -green fluorescent reporter *(Figure 1B, S1A)*. Reporter expression is driven by a cell-type specific GAL4 line. Providing heat shocks at various times is used to reconstruct lineage progression. The morphology of singly-labeled neurons is used to determine neuronal identity (*Figure 1C’’’’*) and counting the number of alternatively-labeled cells determines birth order (*Figure 1C’-C’’’*). To drive ts-MARCM reporter expression, we used *EL- GAL4* that drives in all EL neurons, together with a newly-generated genetic cassette that amplifies GAL4 expression (*Figure S1B*). We find that in segment A1, ELs are produced in the following order: A08x, A08j1, A08j3, A08j2, A08m, A08o, A08c, A08s, A08e3, A08e1, A08e2 (*Figures 1E, S1*). Notably, all these ELs have been suggested to NB3-3A1L/R progeny, and our data provides the first confirmation for several (A08o, A08j1-j3) (Mark et al., 2021)(Wreden et al., 2017)(Heckscher et al., 2015). Because we already knew the first two neurons in the lineage were non-ELs, our data precisely define the birth order for all neurons in NB3-3A1L/R lineage at cellular resolution.

Next, we needed to find all NB3-3A1L/R neurons in the Drosophila L1 larval connectome because this provides high-resolution pictures of neuron morphology including the locations of dendrites (i.e., regions with post-synaptic densities, *Figure 2D cyan*) and axons (i.e., regions with pre-synaptic active zones, *Figure 2D red*). To locate neurons of interest in the connectome we used two criteria. First, lineage-related neurons often have clustered cell bodies and neurites that form a tight bundle entering the neuropile (Schmid et al., 1999)(Mark et al., 2021). We generated a new *NB3-3-GAL4* line, which selectively labels NB3-3 (*Figure S2*). Using both *NB3-3-GAL4* and *EL-GAL4* to drive membrane GFP, we confirmed that the GFP-labeled neurons have clustered soma and bundled neurites (*Figure 2A-B*). Second, we used ts-MARCM data as ground truth to identify neurons with corresponding morphologies in the connectome (*Figure S1*). With these two criteria, we identified a lineage bundle that contained all the ELs plus two other non-EL neurons (*Figure 2C)*. The non-EL neurons were an undifferentiated neuron, which has never been reported before, and a motor neuron, which is consistent with previous reports (i.e., MN 22/23, *Figure 2D top row*)(Schmid et al., 1999). Then, with the high-resolution morphology data, we asked to what extent is EL morphology correlated with birth order and temporal cohort borders. We found one morphological group of early-born ELs, which included A08x, A08j1, A08j3, A08j2, A08m (*Figure 2D second row*). The axons of these ELs project to central brain, with one exception (A08m). Their dendrites are found on both sides of the midline (ipsilateral and contralateral to the soma) and project ventrally from the main neurite, with one exception, A08m, which has only contralateral dendrites. Two of these ELs had been previously identified as early-born ELs (A08x and A08m) (Wreden et al., 2017). The second morphological group contained late-born ELs including A08o, A08c, A08s, A08e3, A08e1, A08e2 (*Figure 2D third and fourth row*). The axons of these ELs project to central brain or remain local. Their dendrites are ipsilateral to the soma and project dorsally from the main neurite, with one exception, A08c, which projects dendrites both dorsally and ventrally. All of these ELs, except A08o, had been previously identified as late-born ELs (Wreden et al., 2017). Thus, we find morphological groupings of neurons in the NB3-3A1L/R lineage that are strongly correlated with neuron birth time and that reflect previous grouping of ELs into early-born and late-born temporal cohorts.

**Figure 2.**
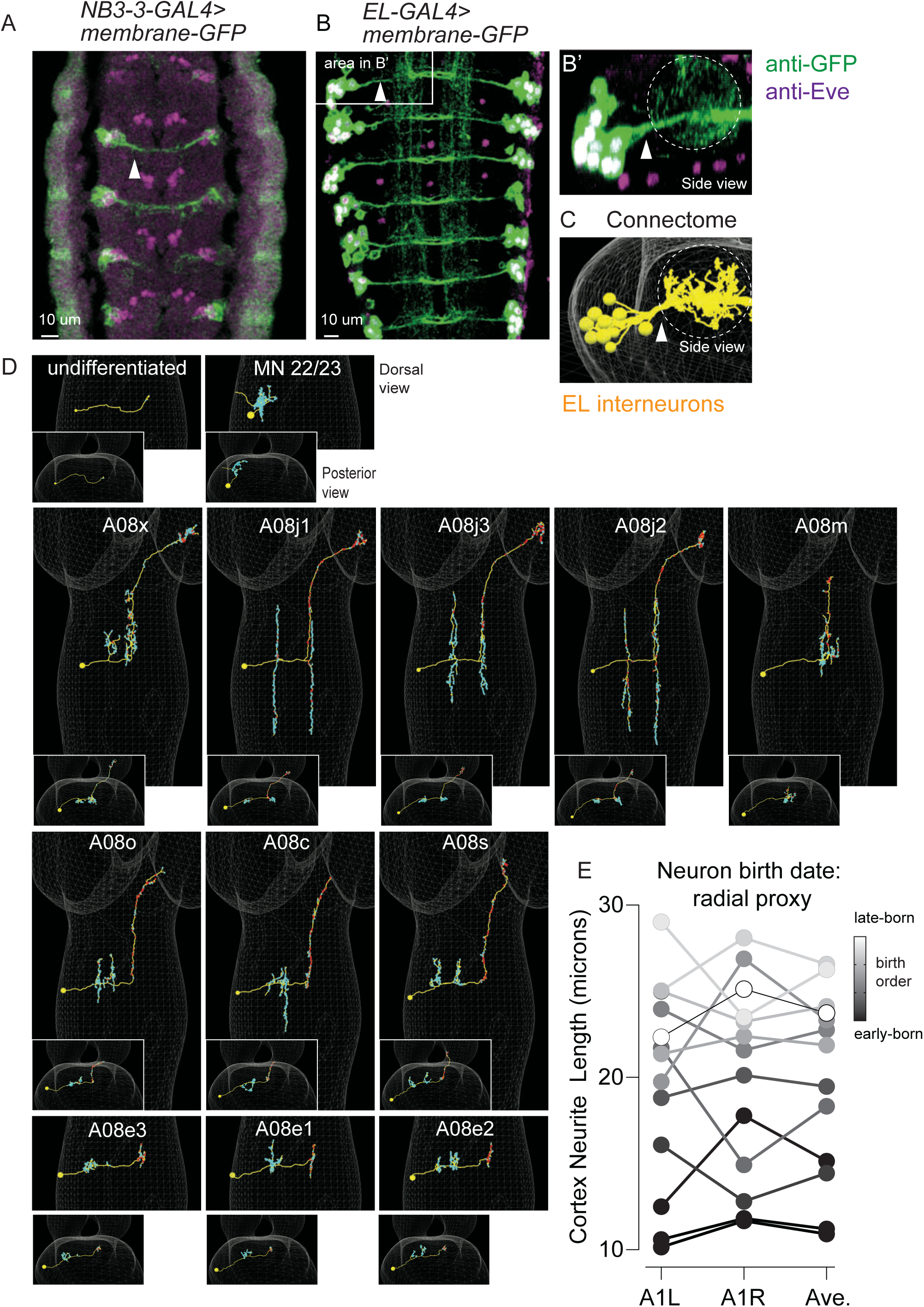
The morphology of all NB3-3A1L/R neurons in the connectome. **A-C. Images of NB3-3A1L neurons with clustered soma and bundled neurites** A-B. *NB3-3-GAL4* or *EL-GAL*4 driving expression of membrane GFP was immortalized using a permanent labeling strategy (Table S4). Arrowheads point to bundles formed by neurons before they enter the neuropile. The images are dorsal views with anterior up. In A, a stage 12 embryo, shows that all NB3-3 neuronal progeny (including two non-ELs) are in a bundle. In B, a first instar larva, shows that in larval stages ELs form a bundle. The box shows segment A1L, which is enlarged and rotated in B’. B’ shows a side view of the bundle as it enters the neuropile (dashes). An image corresponding to B’ in from the connectome is shown in C. **D. Images of each NB3-3A1L neuron in the connectome** For each image, a faint white mesh shows the outline of CNS volume. Large images are dorsal views with anterior up. Smaller images are posterior views (looking towards the brain) with dorsal up. Yellow circles and lines are soma and neurites, respectively. Red and cyan dots are pre- and post-synaptic specializations, respectively. Neuron names are shown at the top of each panel. First-born NB3-3A1L/R progeny are in the top row; early-born ELs in the middle row; and late-born ELs in the bottom two rows. **E. Quantification of NB3-3A1L/R cortex neurite length as a proxy for birth timing** The length of the neurites between the soma and neuropile has been used as a proxy for neuronal birth timing. Plotted on the y-axis are cortex neurite lengths computed for NB3-3 neurons in segment A1L, A1R, and their average. Each dot represents a single neuron (or average of two). Gray scale shows the precise birth order as determined by ts-MARCM. There is a rough correlation between birth order and neurite length, with earlier born-neurons possessing shorter neurites.

In a previous report, Mark and colleagues used the length of a neurite between the soma and the point at which it enters the neuropile as a proxy for neuronal birth order (Mark et al., 2021). This measure is called ‘cortex neurite length’. Here, we sought to quantify the relationship between cortex neurite length and neuron birth order because for the first time we had a complete lineage with both precisely defined birth order information and cortex neurite length data. We plotted cortex neurite length for each neuron in the NB3-3A1L/R lineage, and a found strong positive correlation as measured by Pearson’s correlation (r(9) = 0.82, p= 0.002, and r(9) = 0.77, p = 0.006, for A1L and A1R, respectively)(*Figure 2E*). However, exact birth order cannot be accurately inferred by neurite length. We conclude that it is possible to roughly, but not precisely, infer birth order of neurons within a lineage using only anatomical data.

#### Synaptic inputs onto NB3-3A1L/R neurons

Next, to characterize the relationship between neuronal birth order and patterns of synaptic connectivity, we needed to comprehensively identify all the synaptic inputs to NB3-3A1L/R neurons. To do so, we mined the Drosophila larval connectome, and we identified 4944 synaptic inputs on NB3-3A1L/R neurons, which came from 1179 different neurons (or neuronal fragments) (*Table S1*). We categorized the neuronal sources of NB3-3A1L/R synaptic inputs: a majority (61%) came from other nerve cord interneurons, followed by sensory neurons (19%), with the remainder being from unidentified neurons or neurons of the central brain (*Figure 3A, Table S1*). Because a large majority of inputs onto NB3-3A1L/R neurons come sensory neurons and nerve cord interneurons we focused on these for the rest of this paper.

**Figure 3.**
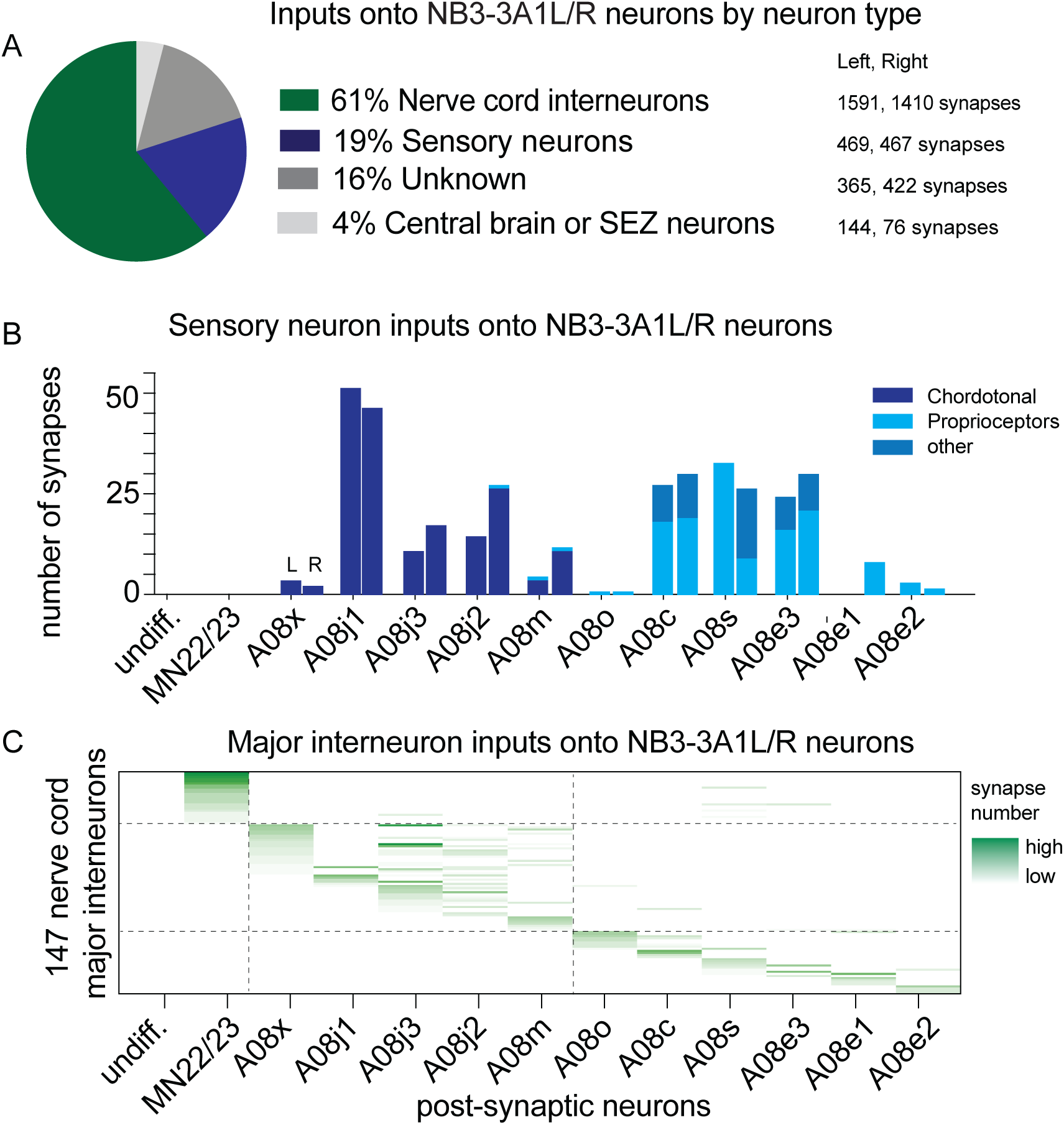
There are sharp transitions in lineage progressing with respect to sensory neuron and interneuron inputs onto NB3-3A1L/R. **A-C. Quantification of synaptic inputs onto NB3-3A1L/R.** A. The pie chart displays the percentage of total synapses from a given neuron type onto NB3- 3A1L neurons. Data are color-coded and the total number of synapses contributed by a given neuron type is shown at right. B. Sensory neurons form synapses onto NB3-3A1L/R neurons. Chordotonal sensory neurons (dark blue) synapse onto early-born ELs. Late-born ELs get input from proprioceptive or other sensory neurons (lighter blues). The X-axis represents the proportion of total synaptic input onto a given neuron that come from a single class of sensory neuron. Rows in the Y-axis represent pairs of post-synaptic NB3-3A1L (L) and NB3-3A1R (R) neurons sorted by birth order. C. Major interneurons form synapses onto NB3-3A1L/R neurons. The X-axis represents 147 nerve cord major interneurons (see Methods). Each column represents one neuron. Names are not shown due to space limitations. Rows in the Y-axis represent post-synaptic NB3-3A1L/R neurons. If an input interneuron forms a synapse with a NB3-3A1L/R neuron, the row-column intersection is shaded green, with darker the green representing greater number of synapses. Dashed lines are placed at the border between non-ELs, early-born ELs, and late-born ELs.

A major functional difference between early-born ELs and late-born ELs is their response to sensory stimuli (i.e., early-born ELs respond to vibration, but late-born ELs do not) (Wreden et al., 2017). So, first, we characterized the relationship between birth order and sensory neuron input. In the NB3-3A1L/R lineage, the earliest born, non-ELs (undifferentiated neuron and MN 22/23) get no input from sensory neurons (*Figure 3B*). Two early-born ELs, A08x and A08m were known to get chordotonal input (Wreden et al., 2017). Here, we newly found that all other early-born ELs get input from chordotonal sensory neurons (*Figure 3B*). We also found that late-born ELs do not get chordotonal input, but instead get input from proprioceptive and other sensory neurons (*Figure 3B*). This agrees with previous finding (Heckscher et al., 2015), and further shows for the first time that the late-born A08o gets a low amount of proprioceptive input (*Figure 3B*). Thus, we conclude that based on input from sensory neurons, there are two sharp transitions in NB3-3A1L/R lineage progression, from non-EL neurons to early-born ELs and from early-born ELs to late-born ELs.

Next, we asked if there were similar, sharp transitions in NB3-3A1L/R lineage progression when looking at inputs from nerve cord interneurons. Note, in the Drosophila larval nerve cord, individual neurons do not contribute equal numbers of synapses onto a given downstream neuron. For example, of the 1179 neurons synapsing onto NB3-3A1L/R neurons, a majority (694) contributed only one or two synapses (*Table S1*). And so here, we analyzed only nerve cord interneurons that provided “major” inputs onto NB3-3A1L/R neurons, or “major interneurons” (see Methods for details). We found 147 major interneurons, which together provided 1990 synapses onto NB3-3A1L/R neurons. In the NB3-3A1L/R, the undifferentiated neuron gets no synaptic input, and MN22/23 gets inputs from major interneurons that largely do not synapse onto any ELs *(Figure 3C*). For early-born ELs, many major interneurons synapse onto multiple neurons within the temporal cohort *(Figure 3C*). Similarly, for late-born ELs, major interneurons synapse onto multiple neurons within the temporal cohort *(Figure 3C*). Importantly, there are nearly no major interneurons that synapse onto both early-born and late-born ELs *(Figure 3C*). We conclude that based on major interneuron input, NB3-3A1L/R lineage progression undergoes sharp transitions in patterns of input connectivity. Further, these transitions are correlated with temporal cohort borders.

The qualitative analysis above suggested that early-born and late-born ELs have distinct connectivity patterns. To quantitatively test this idea, we next analyzed all neuronal input data, not selected subsets, using a network science-based approach, specifically distance analysis. Briefly, we represented the input connectivity for each neuron as a vector. For every pair of neurons, we calculated the Euclidean distance in vector space. This provides a metric for the difference in connectivity between two neurons. Next, we permutated the inputs of each neuron independently, preserving number of inputs, but shuffling input identity. For each shuffled dataset Euclidian distances were computed to provide an estimate of how the same set of neurons could be wired by chance. Real values were normalized to shuffled values to generate a z-score that allowed us to determine statistical significance (see Methods). In examining the distribution of z-scores, we find that early-born ELs have significantly *<u>similar</u>* patterns compared to other neurons in their temporal cohort, but not compared to other neurons in the lineage (*Figure 4A*). Late-born ELs have significantly *<u>similar</u>* patterns compared to other neurons in their cohort, but not compared to other neurons in the lineage (*Figure 4A*). Importantly, in general, early-born and late-born ELs have significantly *<u>different</u>* connectivity from each other (*Figure 4C*). This provides statistical support for the idea that early-born ELs and late-born ELs have sharp transitions in patterns of connectivity beyond what is expected by chance. Further, these sharp transitions in connectivity correlate with previously-characterized circuit function and temporal cohort borders.

**Figure 4.**
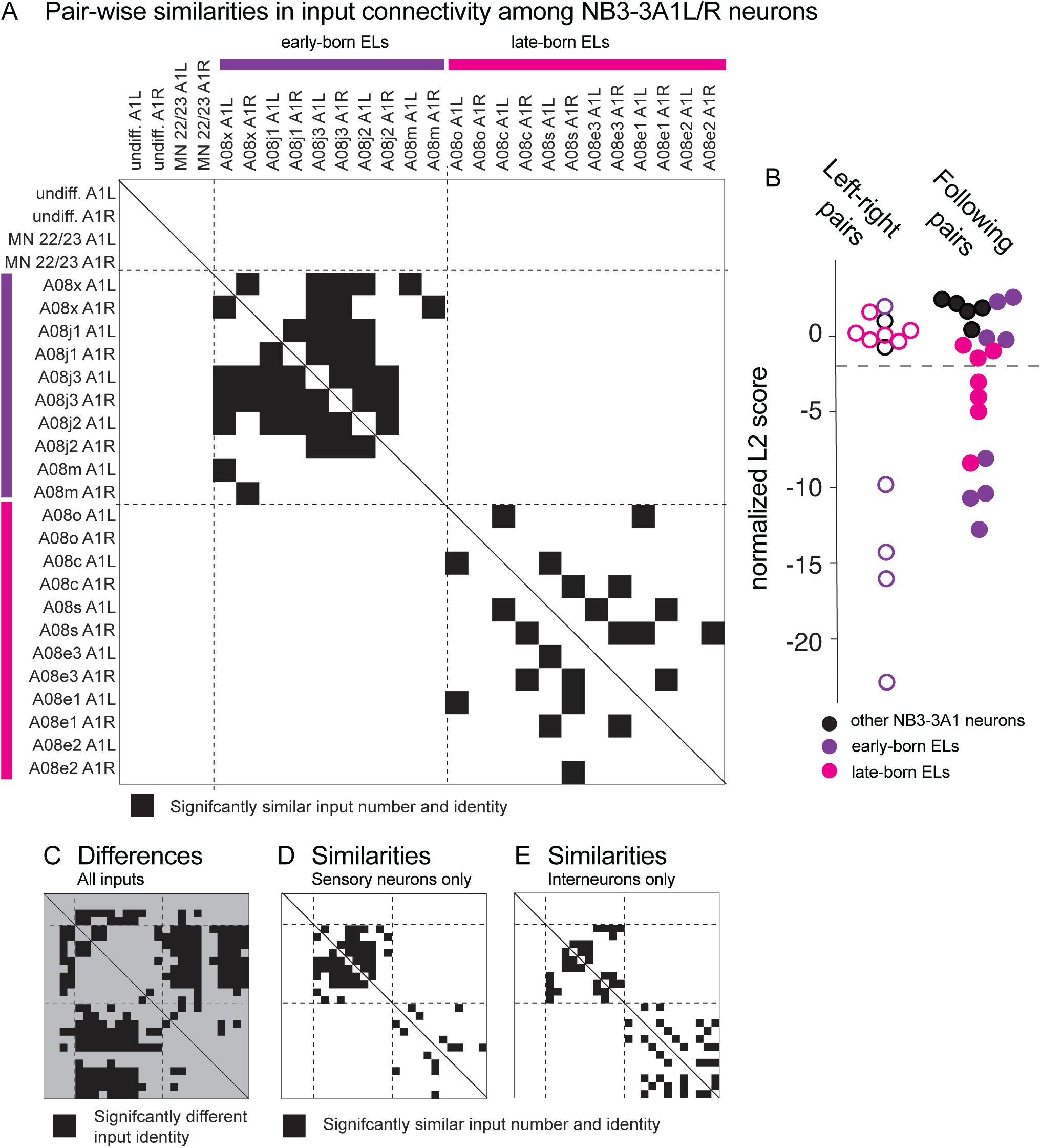
Early-born and late-born ELs have significantly similar input connectivity patterns. **A-E. Quantifications of input connectivity among NB3-3A1L/R neurons using network analysis** A. The plot shows pairwise comparisons of input connectivity with significant similarities in black (normalized, non-binary distance analysis, see Methods). The plot is symmetric, and a solid line shows the diagonal. Left and right NB3-3A1 neurons are arranged in order of their birth on the X-axis and Y-axis. Early-born ELs are indicated with purple and late-born ELs with magenta. Row-column pairs with scores significantly (p<0.05) smaller than shuffled controls are shown in black. Dashed lines are placed at the border between non-ELs, early-born ELs, and late-born ELs. B. The plot shows a summary of pair-wise differences in input connectivity (normalized, binary distance analysis, see Methods). At left is a comparison of left-right neuron pairs (hollow circles) and at right is a comparison of following (in birth order) neuron pairs (solid circles). All left-right pairs with significant similarities are early-born ELs. Both early-born and late-born ELs have significant similarities among following pairs. Significance threshold (p<0.05) is marked in a dashed line. Smaller scores indicate increased input similarity. C. The plot is similar to that in A, but shows normalized Euclidian distance <u>differences</u> between neuron pairs. The background is gray to visually distinguish it from similarity plots. Black marks pairs, whose input identities are significantly different than would be expected by chance. D-E. The plots are similar to that in A, but computed separately with sensory neuron only or interneuron only inputs.

It has been suggested that within a lineage, temporal cohorts could be “copies” of each, implying that although the specific input partners differ, the pattern of connectivity repeats (Wreden et al., 2017). Distance analysis allowed us to asses this idea by helping to visualize connectivity patterns among early-born versus late-born ELs (*Figure 4A*). Early-born ELs had significant similarities between left-right pairs of neurons, whereas late-born ELs did not (*Figure 4B*). Early-born ELs had similarities with many other neurons within the temporal cohort, whereas late-born neurons tend to be most similar to neurons that were born next in the sequence *(Figure 4A-B*). The idea that neurons within a cohort are more similar to neurons next born in the sequence suggests that within a cohort there are graded transitions in connectivity patterns. Ultimately, data that show that temporal cohorts are not merely copies of each other, and they suggest that the computations performed by early-born and late-born ELs differ.

Finally, we used distance analysis to ask what drives the clustering of EL interneurons into two cohorts—connectivity with sensory neurons of PNS, or with interneurons of the CNS, or both? To address this question, we repeated distance analysis, dropping out either interneuron or sensory inputs. For drop out of interneurons, only sensory connections remain (*Figure 4D*). There were still significant similarities in inputs specifically among neurons of a temporal cohort. However, for early-born ELs, the total number of neuron pairs with significant similarities was increased, whereas for late-born ELs, it was reduced. This suggests that interneuron input diversifies the early-born cohort, but unifies the late-born cohort. Data for drop out of sensory neurons is consistent with this idea (*Figure 3E*). This may hint at differing developmental strategies for temporal cohort assembly.

In summary, we find that NB3-3A1L/R lineage undergoes two sharp transitions during lineage progression--from non-EL neurons to early-born ELs and from early-born ELs to late-born ELs. Sharp transitions are seen with respect to morphology and patterns of input connectivity, supporting a sharp transitions model. Further, sharp transitions are well correlated with previously defined early-born and late-born ELs temporal cohort borders (Wreden et al., 2017). Ultimately, these data bring into alignment the concepts of temporal cohorts, circuit function, and circuit anatomy at single neuron resolution.

### Early-born ELs are embedded in a feed-forward circuit and encode the onset of vibrational stimulation

For the rest of this study, we focused on early-born ELs as a model to understand circuit assembly. Early-born ELs and late-born ELs contribute to different circuits (Wreden et al., 2017). In comparison to late-born ELs, the anatomical circuit motif in which early-born ELs are embedded is poorly characterized (Heckscher et al., 2015)(Mark et al., 2021). Stem cell lineage-to-circuit relationships can differ depending on the type of circuit generated (Xu et al., 2014). And so, we needed to characterize the early-born EL circuit.

To understand the circuit to which early-born ELs contribute, we started our analysis by grouping the neurons that synapse onto early-born ELs into classes based on morphology. For this analysis, we used 1945 synapses formed on early-born ELs by 331 neurons (sensory neurons and nerve cord interneurons) (*Table S2*). 6% of all of these synaptic inputs came from Basin interneurons (*Figure 5B*). Basins are a previously-described class of excitatory neurons that have ipsilateral axons and dendrites (*Figure 5D*) (Ohyama et al., 2015b). 12% of synaptic inputs are from A08 interneurons (*Figure 5B*). In experiments described below, we find that A08 interneurons are early-born ELs from segments other than A1. 17% of synaptic inputs come from Ladder interneurons (*Figure 5B*). Ladders are a previously-described group of un-paired inhibitory neurons with cell bodies in the midline and left-right symmetrical arbors (*Figure 5E*) (Jovanic et al., 2016). 31% of synaptic inputs come chordotonal sensory neurons (*Figure 5B*), which we and others have previously shown respond to vibration (Wreden et al., 2017)(Ohyama et al., 2013). Input from these four neuron classes (145 neurons) accounts for 66% of the total nerve cord synaptic inputs. The remaining 34% of inputs come from 186 neurons (*Figure 5E*), a majority of which contribute only one or two synapses onto early-born ELs *(Table S2).* Thus, there is a strong bias in terms of which classes of neurons provide synaptic inputs onto early-born ELs in segment A1.

**Figure 5.**
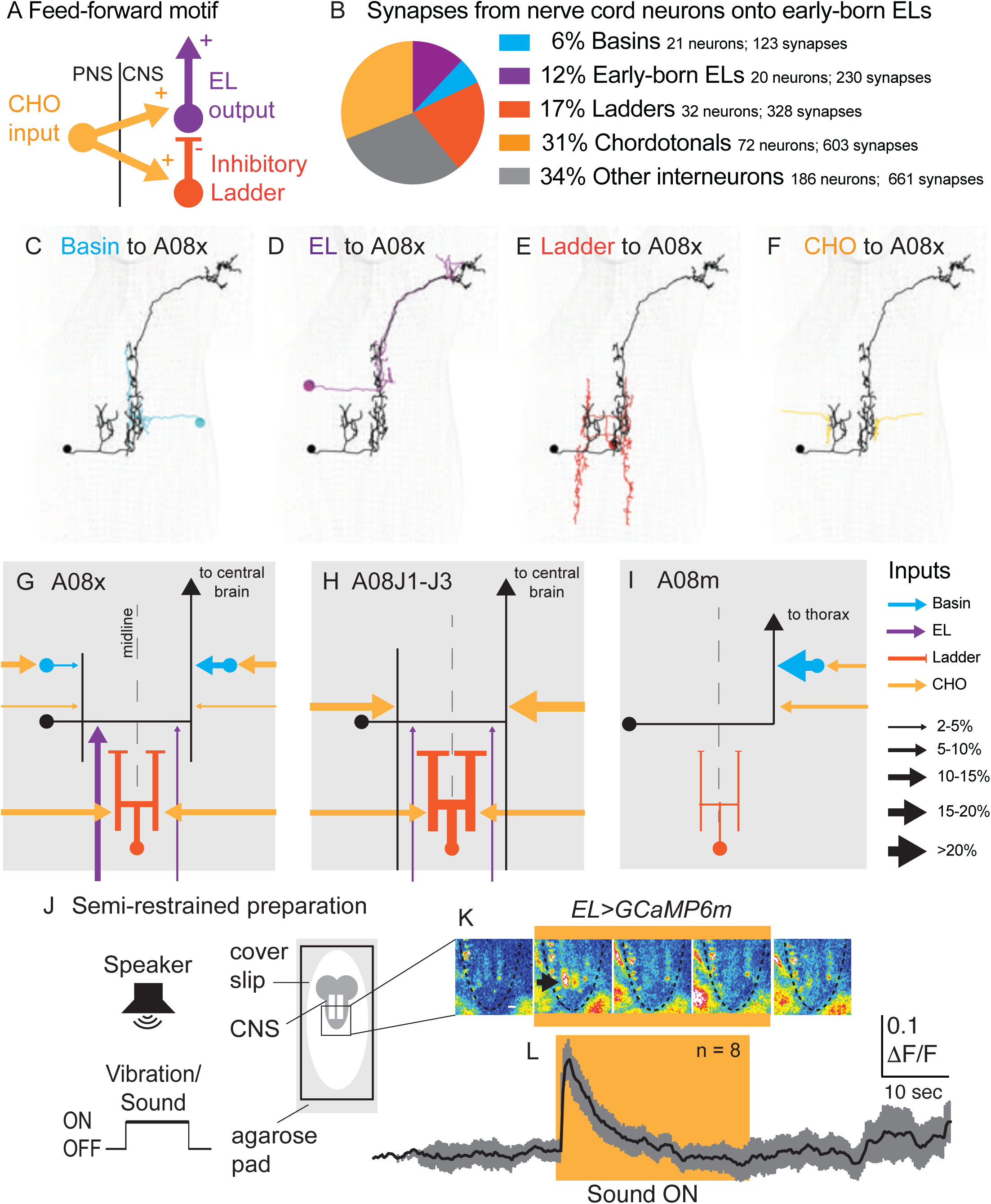
Early-born ELs are embedded in a feed-forward motif and encode the onset of vibrational stimuli. **A. Image of the feed-forward motif in which early-born ELs are embedded.** **B. Quantification of nerve cord synapses onto early-born ELs** The pie chart displays inputs onto early-born ELs. Each slice is the percentage of synapses from a given anatomical class (e.g., Ladders) compared to the nerve cord neuron synapses (Table S2). **C-F. Images of neurons from each class**. The early-born EL, A08x A1L is shown in black. The other neurons with inputs onto A08x are shown in color code in B. **G-I. Illustrations of patterns of synaptic inputs onto different early-born ELs** ELs are in black. Arrows are excitatory connections and bars are inhibitory (i.e., Ladders). The key at right shows how line thickness corresponds to percentage of inputs onto a given neuron. **J. Illustration of the preparation and stimulus protocol used for calcium imaging** A larva is placed on a bed of agarose with a cover slip on top. Fluorescence in the CNS (gray lobed structure with two white lines [neuropile]) is recorded before, during, and after a sound is played from a speaker. **K. Images from representative recordings of calcium signals in ELs.** Frames from a representative recording are shown at 8 second intervals. Yellow box indicates frames where sound stimulus was ON. Images are pseudo-colored with white/red as high fluorescence intensity and blue as low. Anterior is up. Scale bar is 50 microns. Dashed lines show the outline of the nerve cord. The black arrow in the second image panel indicates a region of neuropile with increased fluorescence. **L. Quantifications of EL calcium signals** Changes in EL calcium signaling upon vibrational stimulus (yellow box) show a rapid increase followed by decay. The black line represents average fluorescence intensity and gray represents standard deviation. ΔF/F is the change in fluorescence over baseline. N = number of larvae recorded. For genotype see Table S4.

After identifying these four classes of inputs, we next asked how they were anatomically arranged at the level of circuit motif. Here, we consider early-born ELs to be the circuit outputs because they are excitatory and project to the central brain (or, for A08m, thorax) (*Figure 5G-I*). Input to the circuit comes from chordotonal sensory neurons. Chordotonal sensory neurons provide <u>direct</u> excitatory input onto early-born ELs (yellow arrows*, 5G-I*), and they provide indirect <u>input</u> excitatory and inhibitory onto early-born EL in A1 via Ladders, Basins and other early-born ELs (orange, blue, and purple arrows respectively, *Figure 5G-I*). Thus, anatomically, early-born ELs in A1 are imbedded in a feed-forward circuit motif.

Because early-born ELs get direct excitatory and indirect inhibitory input from chordotonal sensory neurons (*Figure 5A*), this suggested early-born ELs could be activated upon initial chordotonal stimulation and inactivated with delay. Our previous calcium imaging experiments used a slow, pre-synaptic calcium sensor, and showed early-born ELs respond to chordotonal stimulation (Wreden et al., 2017). Here, to get a more precise understanding of dynamics of EL activity, we monitored stimulus encoding using a cytoplasmic calcium sensor with faster signal decay. In response to vibration, which activates chordotonal sensory neurons (Wreden et al., 2017), EL activity initially peaks and then the signal rapidly declines, although the vibrational stimulus remains (*Figure 5J-L)*. These data support the idea that functionally at least a subset of ELs encode stimulus onset.

In summary, we find that all early-born ELs are embedded in a feed-forward circuit motif and that at least a subset of ELs encode the onset of vibrational stimuli.

### Highly selective wiring among temporal cohorts

After characterizing the circuit containing early-born ELs, we were in the position to test the hypothesis that this feed-forward circuit is assembled by preferential connectivity between distinct temporal cohorts (Meng & Heckscher, 2021). To test this, we needed to know to what extent could Basin, Ladder, and A08 interneurons be considered temporal cohorts. To be a temporal cohort, interneurons must be lineage-related and born within a tight time window. Here, we took a combination of light microscopy and connectome-based approaches to address this question.

#### Basins are a middle-to-late-born temporal cohort from NB3-5

There are four pairs of Basin interneurons per segment, Basin 1-4 (Ohyama et al., 2015b)(Jovanic et al., 2016). Basins have been suggested to be a temporal cohort from NB3-5 (Wreden et al., 2017). NB3-5 is one of the first neuroblasts to form in Drosophila embryos (J. J. Broadus et al., 1995). It is among the largest embryonic lineages, reportedly producing up to 36 neurons, a subset of which are found at larval stages (*Figure 6A*) (Monedero Cobeta et al., 2017) (Moris-Sanz et al., 2014). The first-born neurons express the CCAP neuropeptide, and many NB3-5 neurons die during embryogenesis (Monedero Cobeta et al., 2017) (Moris-Sanz et al., 2014). Otherwise, the cell types produced by NB3-5 are poorly understood.

**Figure 6.**
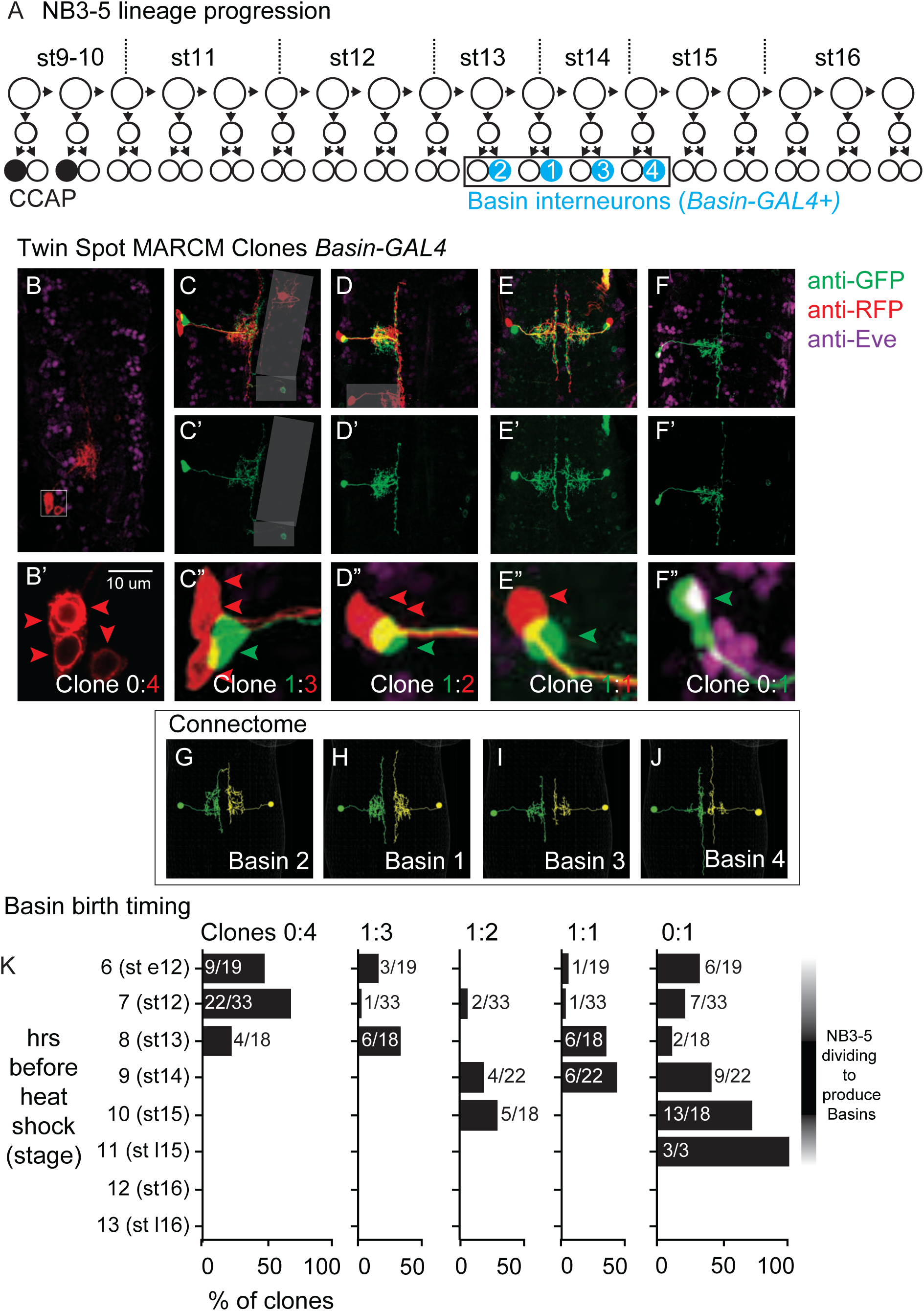
Birth time and birth order of Basin interneurons using ts-MARCM. **A. Illustration of NB3-5 lineage progression** Each circle represents one cell, and each arrow represents a cell division. The X-axis represents developmental time, and dashes represent approximate positions of embryonic stages (e.g., st16). NB3-5 generates two CCAP(+) neurons (black) followed by a series of other neurons. Basins (cyan) are born a middle-to-late window. **B-J. Images of Basin ts-MARCM clones and Basins in the connectome** Examples of ts-MARCM clones are shown in B-F’. Eve staining serves as a counter-stain to visualize the nerve cord architecture. RFP and GFP show ts-MARCM Basin clones. Genotype in Table S4. B shows an example of a clone produced by an early, brief heat shock (see Methods). In this nerve cord, only one clone is present, and in that clone, all four Basins are labelled. B’ corresponds to the boxed region in B. Red arrowheads point to each cell body (two are stacked on each other). C-F” show examples of other clone types. C-F shows two color labeling. C’-F’ shows morphology of the singly labeled Basin. C”-F” show higher magnification of the cell bodies with arrowheads pointing to labeled cells. Clone types indicated at the bottom of the panel. G-J show left-right pairs of Basin interneurons in segment A1 of the connectome. All images shown in dorsal view with anterior up. **K. Quantification of types of clones produced by variously timed heat shocks.** The clone type is displayed at the top of each graph. Y-axis for each graph shows the various times after egg collection until heat shock was applied. The X-axis for each graph represents the percentage of clones of that clone type that were produced by a given heat shock protocol. Numbers are the total number of clones of a clone type over the total number of clones scored for that time point.

First, we asked if Basins are likely to be a lineage-related set. To do so, we used ts-MARCM, and we detected ts-MARCM recombination events using *Basin-GAL4*, which is expressed in all Basin interneurons (Ohyama et al., 2015b). First, we induced ts-MARCM clones using brief heat-shocks at early times in development. This produced one or two clones per CNS (e.g., *Figure 6B*) consistent with the idea that recombination levels were low. We counted the number of Basin neurons in each clone and found that most often all four Basins were labeled (e.g., *Figure 6B’, K).* This is consistent with the idea that recombination occurred in a single dividing neuroblast and that neuroblast produces all Basin interneurons. Next, we used ts-MARCM to determine the relative birth timing of Basins within the lineage. In this set of experiments, we provided heat shocks at a variety of developmental time points and scored the resulting progeny. We find that Basins are born in the following order: Basin 2,1,3,4 (*Figure 6A-J*). Additionally, based on the types of clones produced at different times (see Methods for details), our data suggest Basins are born within a middle-to-late window during NB3-5 lineage progression (*Figure 6K*). We conclude Basins are a lineage-related set of neurons born within a tight time window, consistent with the idea that Basins are a temporal cohort.

Next, we wanted to directly test the idea that Basins are progeny of NB3-5. To do so, we crossed a fly line expressing membrane-GFP in all Basins (*Figure 7A)* to three different fly lines, each expressing nuclear localized (nls) RFP in the NB3-5 lineage neurons. In all cases, we observed co-expression of Basin and NB3-5 lineage markers (*Figure 7B-D’*). We conclude that Basins are progeny of NB3-5.

**Figure 7.**
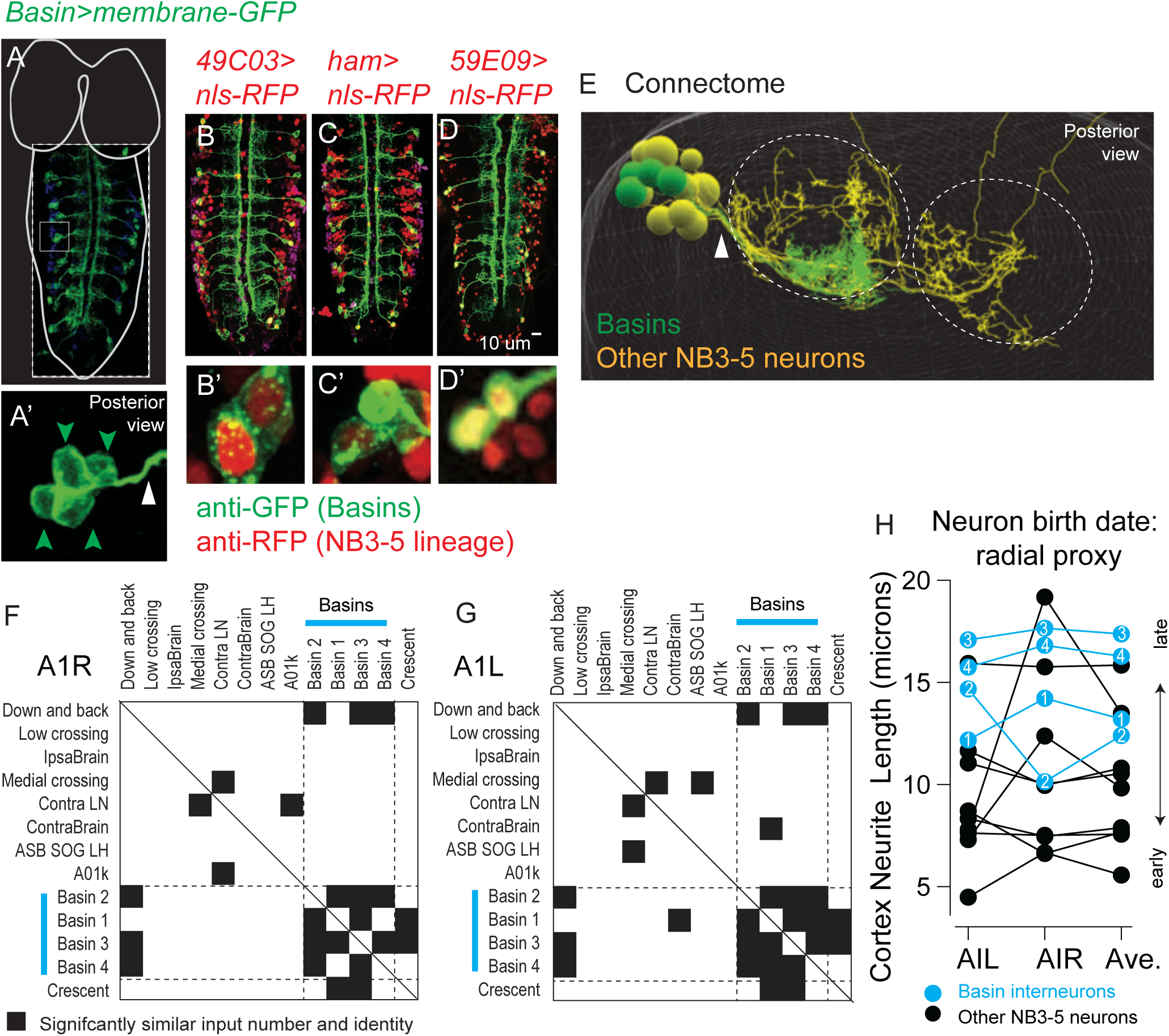
Basins are a middle-to-late born temporal cohort in the NB3-5 lineage. **A-E. Images of Basin interneurons and other lineage-related neurons** A. The larval CNS with Basins neurons expressing membrane GFP is shown in dorsal view with anterior up. For context, the outline of the nerve cord and brain lobes is shown in white. Dashed box outlines the image. The small light box region is shown in A’. The green arrowheads point to Basin cell bodies, which are clustered. The white arrow points to bundle of Basin neurites. B-D’ Larval nerve cords show co-expression of a Basin marker (membrane GFP) and various NB3-5 progeny markers (nuclear localized RFP). Genotypes in Table S4. E. An image of NB3-5 progeny in the connectome is shown. The neuropile is outlined by dashed circles. Basin neurons are in green, other neurons in the NB3-5 lineage are in yellow. Arrowhead points to the bundle containing Basins. **F-H. Quantification of Basin features using connectome data** F-G. To quantify the similarity in wiring among neurons in the NB3-5 lineage normalized (non-binary) distance plots were generated. Right and left NB3-5A1 neurons are arranged in approximate order of their birth (based on average cortex neurite length) with Basins indicated by cyan bars. Row-column pairs with scores significantly (p<0.05) smaller than shuffled controls are in black. The distance analysis plot is symmetric (solid line for the diagonal). Dashed lines are placed at the border between Basins and non-Basins. H. The approximate the birth order of Basins with in the NB3-5 lineage was determined using cortex neurite length as a proxy. The Y-axis plots cortex neurite lengths for NB3-5 neurons in segment A1L, A1R, and their average. Compared to other neurons in the NB3-5 lineage Basins are born near the end of the lineage.

Basin neurons had already been identified in the connectome (Ohyama et al., 2015a). And so, next, we used connectome analysis as an independent means of characterizing the developmental of the Basins. First, we observe Basin cell bodies are clustered and their neurites form a bundle both in light microscopy images and in the connectome (*Figure 7A, E*). This provides further support for the idea that Basins are a lineage-related set of neurons. Second, we assayed Basin birth order using the length of cortex neurite length as a proxy. To do so, in connectome, we identified all neurons in the lineage bundle that contained Basin neurons (*Figure 7E, yellow*) because these neurons are likely to be part of the NB3-5 lineage. Then we measured and plotted the neurite lengths of putative NB3-5 neurons in the hemilineage that included Basins. In this set, Basins possess neurites that are among the longest (*Figure 7H*). These data are consistent with our ts-MARCM data, which suggest that Basins are middle-to-late born. Therefore, by two independent means we find that Bains are a lineage-related set of neurons born within a tight time window. This provides strong evidence that Basins are a temporal cohort.

In the first section of this paper, we found that among neurons in the NB3-3A1L/R lineage there are sharp transitions in input connectivity patterns correlated with temporal cohort boundaries. However, it remained unclear if this was true for early-born and late-born temporal cohorts only, or if this represented a more general “rule”. Because we identified Basins as a temporal cohort, we next examine the relationship between patterns of connectivity and Basin temporal cohort borders. As described in the first section of this paper, using distance analysis, we quantified the significant similarities in neuronal input, this time analyzing neurons in the Basin hemilineage (*Table S3*). We find Basins have statistically significant similarities in input connectivity with themselves compared to most other NB3-5 neurons (*Figure 6G-H*). Thus, there are sharp transition in connectivity patterns correlated with Basin temporal cohort borders (see Discussion for more).

#### Ladders are a late-born temporal cohort from MNB

Ladders have been suggested to be progeny of MNB (Babski et al., 2019). MNB, or midline neuroblast is the only unpaired neuroblast (Wheeler et al., 2009). It produces up to three Ventral Unpaired Midline motor neurons before generating up to six GABAergic unpaired midline interneurons (*Figure 8A*)(Schmid et al., 1999). In the A1 segment of the connectome, there are six unpaired Ladder interneurons with neurites that enter the neuropile in a single tight bundle (*Figure 8B*). We confirmed Ladders are progeny of MNB by crossing a fly line expressing membrane-GFP in one Ladder to three different fly lines, each expressing nuclear localized (nls) RFP in the MNB lineage. In all cases, Ladder and MNB lineage markers are found in the same cell (*Figure 8C-E*). In addition, we used distance analysis to evaluate input connectivity among Ladders in the A1 segment (*Table S3*), and find statistically significant similarities (*Figure 8F*). We conclude that Ladders are a late-born temporal cohort from MNB. Like other temporal cohorts, we find the Ladder temporal cohort border correlates with sharp transitions in synaptic connectivity.

**Figure 8.**
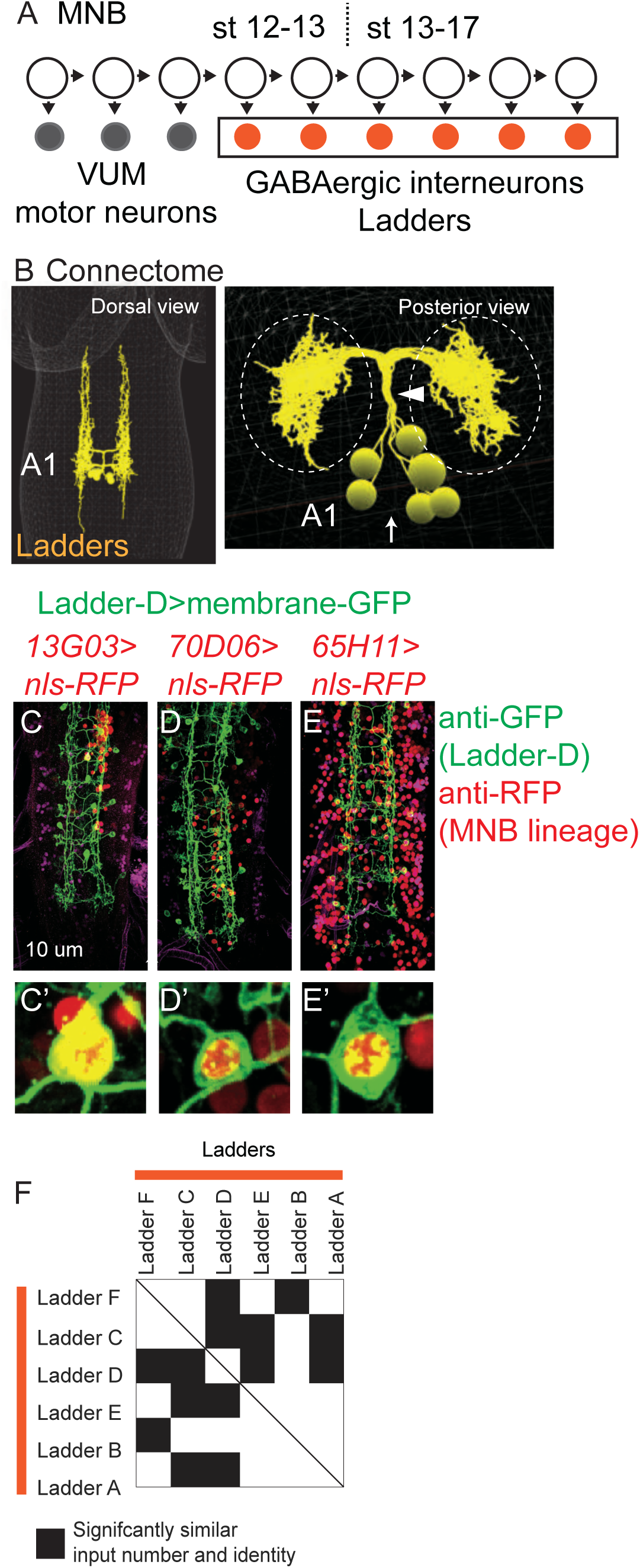
Highly selective wiring among temporal cohorts. **A. Illustration of MNB lineage progression.** Each circle represents one cell, and arrows represent cell division. The X-axis represents developmental time, and dashes represent approximate positions of different embryonic stages (e.g., st16). MNB generates up to three, early-born Ventral Unpaired Midline motor neurons (gray) followed by a series of GABAergic interneurons (orange). These interneurons are Ladders. **B-E. Image of Ladder interneurons in the connectome and of co-expression of Ladder-D interneuron and MNB lineage markers** B. Ladders in segment A1 of the connectome shown in dorsal view and side views. Ladders form a bundle (arrowhead) before entering the neuropile (dashed circles). Midline shown with arrow. C-E’. First instar larval CNSs are shown in ventral view with anterior up. In insets notice co-expression of Ladder-D membrane marker (green), with nuclear localized MNB lineage maker (red). Genotypes in Table S4. **F. Quantification of statistical similarities in Ladder synaptic inputs** Plot show normalized (non-binary) Euclidean distance similarity between pairs Ladders neurons. Ladders are arranged in approximate order of their birth based on cortex neurite length. Row-column pairs with scores significantly (p<0.05) smaller than shuffled controls preserving the inputs number and magnitude are in black. Euclidean distance plot is symmetric (solid line for the diagonal).

#### A08 neurons that synapse with early-born ELs in A1 are early-born ELs from other segments

We noticed that at least some A08 interneurons that synapses with early-born ELs in A1 resembled early-born ELs in terms of morphology. Therefore, we tested the idea that these A08 interneurons could be NB3-3 progeny in segments other than A1. To do so, we used the Multi Color FLP Out (MCFO) system to highlight the morphology of individual neurons within the expression domain of a GAL4 line (Nern et al., 2015). For GAL4 lines, we used either *early-born EL-GAL4* or *late-born EL-GAL4* (Wreden et al., 2017). We compared early-born and late-born EL morphologies to the A08 interneurons that formed synapses with early-born EL in A1. We found a majority were early-born ELs from segments other than A1, including anterior (i.e., thorax) and posterior (i.e., A2) segments (*Figure 9A-H, Table S1*). This reveals intersegmental synaptic connectivity among early-born ELs. Functionally, ELs are intersegmentally coordinated (Heckscher et al., 2005). The observed intersegmental synaptic connectivity could help explain this observation.

**Figure 9.**
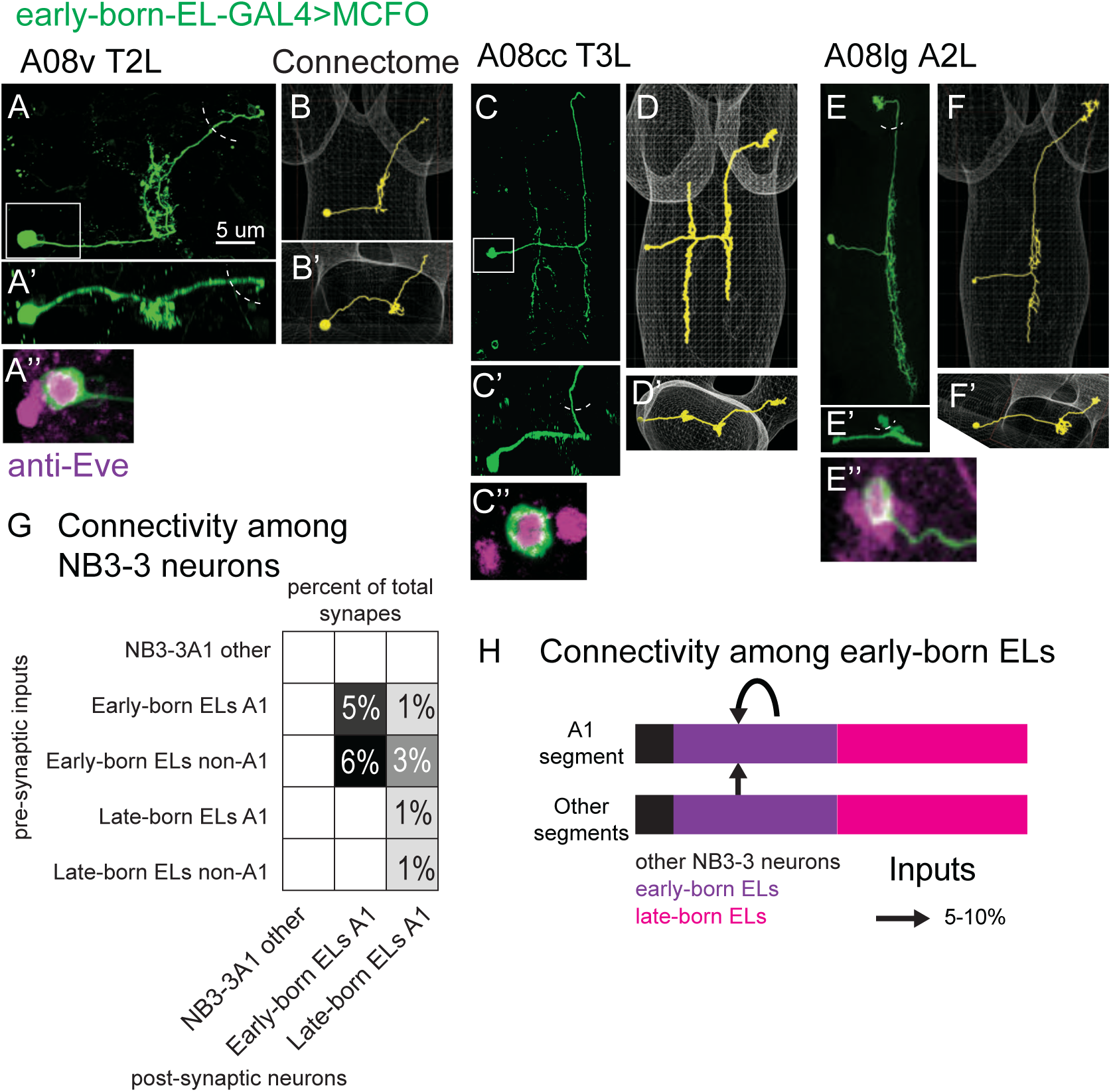
A08 interneurons that synapse onto early-born ELs in A1 are early-born ELs from segments other than A1. **A-F. Images of early-born EL interneuron MCFO clones along with matching A08 interneurons in the connectome** For each figure panel, the main image (e.g., A) shows neuronal membrane labeling in a dorsal view with anterior up. The same cell is shown in side view (e.g., A’). Boxed areas are magnified to show soma co-labeling with Eve, which demonstrates it is an EL (e.g., A’’). Dashed semi-circles shows approximate position of central brain lobes. Confocal images (e.g., A) are shown adjacent to the matching neuron in connectome (e.g. B). Genotype in Table S4. **G. Quantification of intersegmental connectivity among early-born EL interneurons** Each row is a group of pre-synaptic neurons (e.g., Early-born ELs in segment A1). Each column is a group of post-synaptic neurons. White boxes at intersections indicate no connectivity among two groups. Maximal interconnectivity comes from early-born ELs from segments other than A1 synapsing onto early-born ELs in A1 (e.g., 6% of total inputs), followed by connectivity among early-born ELs in A1. **H. Illustration of connectivity among early-born ELs** Bars represent NB3-3 neurons, with black being non-ELs, purple early-born ELs, and magenta late-born ELs. There are synapses among early-born ELs (black arrows).

Because we found intersegmental synaptic connectivity among early-born EL temporal cohorts, we wondered if this occurred among other temporal cohorts. We found intersegmental connections among Basins and Ladders, but not late-born ELs (*Figure 9G-H, Tables S1, S3).* In addition, in A1, we found within-segment (intra-segmental) synaptic connections among early-born ELs, Basins, Ladders, but not late-born ELs (*Figure 9G-H, Tables S1, S3).* One commonality between early-born, Basins, Ladders, but not late-born ELs, is that they receive synapses from chordotonal sensory neurons. This raises the possibility that inter- and intra-segmental synaptic connectivity among neurons in these temporal cohort is a feature important for the processing of vibrational stimuli. Further, these data reveal differences in connectivity among temporal cohorts, hinting at complexity in mechanisms driving their wiring.

In summary, we find Basins neurons are a mid-to-late-born temporal cohort from NB3-5. Ladders are late-born temporal cohort from MNB. And A08 interneurons are early-born ELs from segment other than A1. This supports the hypothesis that in the Drosophila larval nerve cord circuits are assembled by preferential wiring among temporal cohorts. Further, we note that temporal cohorts that form synapses on early-born ELs come from just three of the 30 neuroblast in the nerve cord, demonstrating high selectivity.

### Pre-synaptic interneurons are born after their post-synaptic interneuron partners

To more fully understand assembly of this feed-forward circuit motif, we needed to understand the relative birth timings of early-born ELs, Basin, and Ladder interneurons. A prominent model for spinal cord circuit development suggests that early-born neurons wire with each other to generate circuits driving fast movements, whereas later-born neurons wire with each other to generate circuits driving refined movement (i.e., early-early and late-late model) (Fetcho & McLean, 2010). Thus, here, we test the hypothesis that early-born ELs wire with other neurons born at the same time. Note that in the experiments described above, we determined birth-order of neurons within a given lineage, but had no information about the relative birth timing of neurons between different lineages.

To determine the relative birth timing of early-born Els, Basins, and Ladders we started by asking when during neurogenesis each EL could be generated. We used ts-MARCM with *EL-GAL4.* We supplied heat shocks at different time points, and scored the identity of singly labeled neurons. Heat shock supplied after 7 hours of development labeled a mixture of early-born and late-born ELs, whereas heat shock supplied after 9 hours labeled exclusively late-born ELs (*Figure 10A-B*). Thus, heat shocks after ∼9 hours were not sufficient to label early-born ELs. Next, we asked when during neurogenesis the Basins and Ladders are generated. We used ts-MARCM with GAL4 lines that labeled individual Basin or Ladder neurons (Basin-1 or Ladder-D)(*Figure 10C-D*). We supplied heat shocks at different time points and scored for the proportion of CNS in which clones were induced. In both cases, heat shocks provided at 11 hours of development, or later, failed to label these neurons. Whereas heat shocks provided at 10 hours of development, or earlier, resulted in labeling (*Figure 10E*). Thus, heat shocks at ∼10 hours are sufficient to label Basins and Ladders. Taken together these data strongly suggest that Basins and Ladders are generated after the early-born ELs. Notably, this is inconsistent with the model in the field.

**Figure 10.**
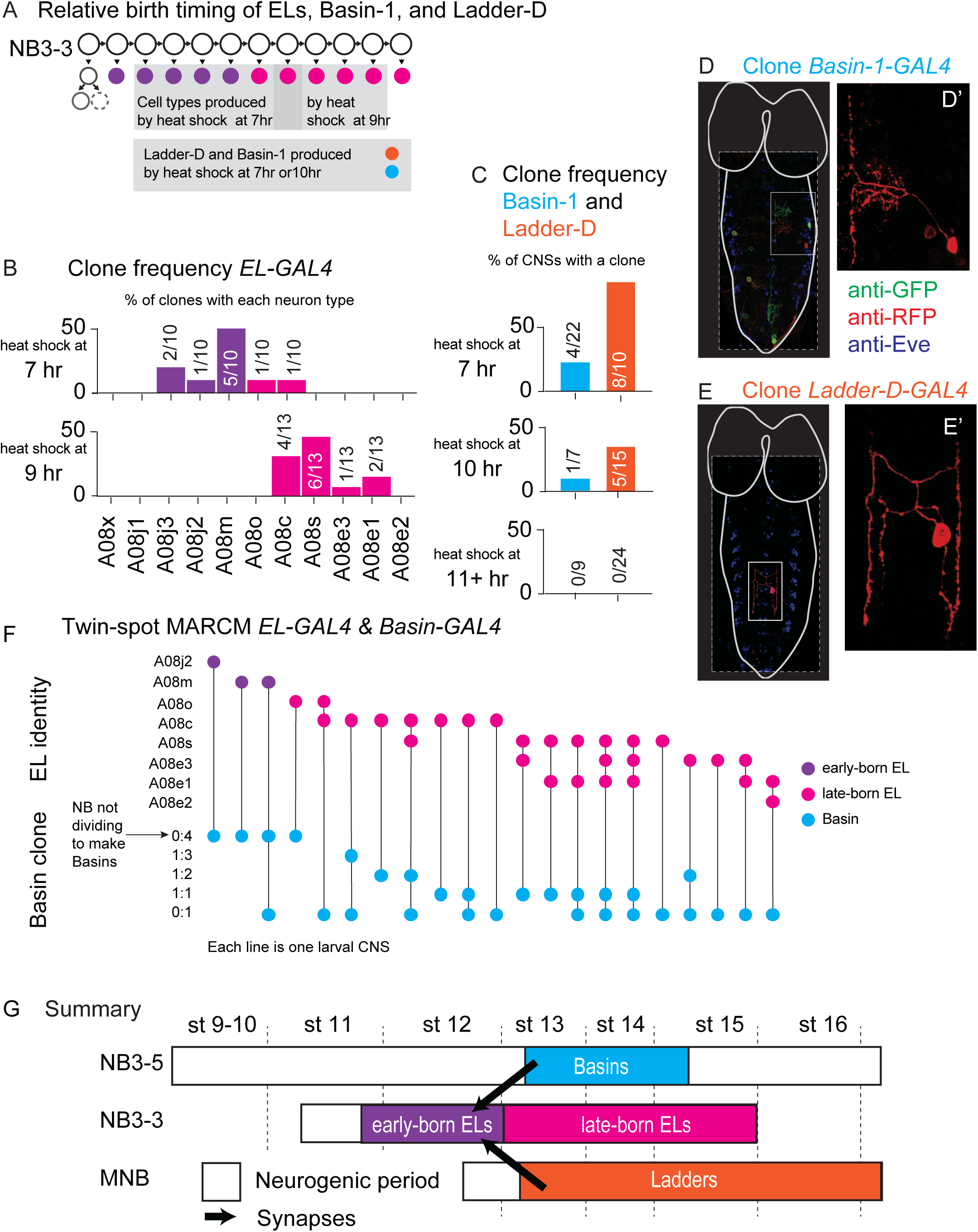
Basin and Ladder interneurons are born after early-born EL interneurons. **A. Illustration of neurons labeled by various heat shock protocols** At top, NB3-3 lineage progression is shown. Each circle represents one cell, and each arrow represents a cell division. Early-born ELs (purple) are labeled during a heat shock at 7 hrs after egg collection (top left gray box). Most late-born ELs (magenta) are labeled during a heat shock at 9 hrs after egg collection (top right gray box). Heat shocks provided at both 7 and10 hrs after egg collection are sufficient to label Basin-1 and Ladder-D (bottom gray box) **B-C. Quantification of cell types labeled by various heat shock protocols** B-C. For each graph, the X-axis represents the type of neuron produced. For ELs (in B) neuron names are presented in birth order. The Y-axis represents how often a given neuron was generated by that protocol with early-born ELs in purple, late-born ELs in magenta, Basin-1 in cyan and Ladder-D in orange. In B, numbers in each bar represent the number of labeled neurons of each type over the total number of neurons scored. In C, numbers in each bar represent the number of CNSs with a labeled neuron over the total number of animal CNSs scored. Genotype in Table S4. **D-E. Images of singly-labeled Basins or Ladders** Larval CNSs are shown in ventral view with anterior up. For context, the outline of the nerve cord and brain lobes is shown in white. Dashed box outlines the image. The small light box shows region in D’-E’. D’ shows a singly-labeled Basin-1. E’ shows a singly-labeled Ladder-D. Genotype in Table S4. **F. Quantification of ELs and Basins labeled in a single CNS.** The Y-axis is divided into two sections. The top shows EL identity and describes a specific A1 EL interneuron type (e.g., A08j2) listed in order of their birth. The bottom shows Basin clone type and refers to the pattern of labeling of Basin neurons. For example, 0:4 means four neurons were labeled in one color and none another. Each column (X-axis) represents a different larval CNS, with a dot indicating the type of neuron labeling observed. A line connecting the two clone types to help visualize the pairs. In some CNSs, more than one type of Basin clone was produced, and so multiple dots are present. **G. Illustration of relative birth timing of Basins, Ladders, and early-born EL interneurons.** Each row represents neurons generated from a different neuroblast over time. The X-axis represents developmental time, and dashes represent approximate positions of different embryonic stages (e.g., st16). Early-born ELs (magenta) get input from neurons born after they are born.

Because our data were inconsistent with such a prominent model, we performed an additional set of experiments to confirm our findings. Specifically, we performed dual ts-MARCM using both *Basin-GAL4* and *EL-GAL4* in the same larva. This approach allows for direct comparison of neurons generated at a given time in a single animal. We provided a series of heat shock and scored the resulting clones as follows: EL clones were scored for neuron identity (e.g., A08j2), and this was used to assign ELs as early-born or late-born. Basin clones were scored for clone type (e.g., 1:3 or 1 neuron labeled in one color, 3 neuron in another color) and this was used to determine the time window of Basin production. Specifically, if all Basins were labeled in a single color (0:4 clone), the heat shock was provided before the neuroblast began to divide to produce Basins. Alternatively, if Basins were singly-labeled (e.g. 1:3, 1:2, etc), the heat shock was provided to cells as they were dividing to produce Basins. Within our dataset, three animals, had labeled early-born ELs. In all of these animals, Basins were labeled in a single color, consistent with the idea that the neuroblast had not yet begun generating Basins (*Figure 9F*). In 17 animals, we found singly-labeled Basin neurons, suggesting Basins were being generated. In 16 of these 17 animals, late-born ELs were labeled (*Figure 9F*). Together these data provide strong evidence that early-born ELs are born earlier than the Basin interneurons.

In summary, we find that among neurons in the feed-forward circuit, early-born EL are born before Basin and Ladder interneurons (*Figure 10G*). More generally, this demonstrates that circuit outputs are born before circuit inputs. Further, this rules out the strict (early-early late-late) hypothesis that neurons born at the same time are circuit partners.

## Discussion

In this study we addressed two questions about the stem-cell based assembly of neuronal circuits. First, what is the relationship between neuronal birth order within a lineage and patterns of synaptic connectivity at the single neuron level? Second, how do neurons from different lineages wire with each other? We characterized the birth order, morphology, and input connectivity of all neurons in the NB3-3A1L/R lineage at single neuron and single synapse resolution (*Figures 1-4, S1-S2, Table S1*). We identified a feed-forward circuit that encodes the onset of vibrational stimuli *(Figure 5).* And, for a majority of nerve cord interneurons within this circuit, we identified their stem cell parent, birth order within their lineage, and birth timing relative to each other (*Figures 5-9, Table S2-4*). Together this identifies four temporal cohorts, all of which have sharp connectivity boundaries. Further, it shows that in the Drosophila larval never cord, temporal cohorts associate with each other in specific ways. Instead of following the vertebrate model of early-early and late-late connectivity, temporal cohorts assemble sequentially, with circuit output neurons born before circuit input neurons. Ultimately, this study provides new tools and new insights into the stem cell-based assembly of a fundamental circuit motif.

### New resources for analysis of connectome data

The Drosophila larval connectome is a resource that can be used to understand circuit assembly (Meng & Heckscher, 2021). However, because the connectome is an anatomical dataset, a major challenge is to develop approaches that connect anatomy to developmental origin. In this paper, we develop several approaches. For example, we optimized ts-MARCM for use in Drosophila embryos (*Figures 1, S1, 6, 10*). ts-MARCM is the gold-standard for determining birth order in Drosophila (H.-H. Yu et al., 2009). However, for technical reasons, ts-MARCM has been used only in adults. We discovered ts-MARCM clones can be robustly generated in early stage larvae with the addition of an amplifying and immortalizing gene cassette (*Figure S1B*). In addition, we independently validated several recently-developed methods for inferring developmental origin based on anatomical features in the Drosophila larval connectome (Mark et al., 2021). Specifically, our validations include use of neurite bundles as a proxy for lineage-relatedness (*Figures 2B-C, 7A,E, 8B*) and use of cortex neurite length as a rough proxy for birth order within a lineage (*Figures 2E, 7F*). Finally, we adapted network-science based methods (distance analysis) to characterize the patterns of connectivity in connectome data (*Figures 4, 7, 8*). These approaches should be useful for Drosophila neurobiologists and beyond.

We also developed NB3-3A1L/R as the first entire lineage for which birth order, morphology, and input connectivity is known at single neuron and synapse precision (*Figures 1-4, S1, Table S1*). One reason this is important is because NB3-3 has been extensively studied in embryos and much is known about molecular marker expression of NB3-3 progeny at the single neuron level (Tsuji et al., 2008)(Wreden et al., 2017)(Baumgardt et al., 2014). Because our dataset achieves cellular resolution, good guesses about the embryonic molecular-larval morphological pairings can be made using single neuron birth order as a point of cross reference. Such integrated data generates detailed and testable predictions. For example, our data predict that Castor expression in the late-born ELs promotes projection neuron morphology. Additionally, here, we generated a new *NB3-3-GAL4* line that can be used manipulate gene expression in NB3-3 (*Figure S2*). Thus, NB3-3A1L/R is a model lineage in which transcription factor expression in neuronal stem cells can be linked to circuit assembly and tested for function, and we have generated tools that will enable hypothesis testing in the future.

### Early-born ELs are embedded in a potentially-conserved, feed-forward circuit motif

A circuit motif is pattern of synaptic connections between a set of specific neuron types that can be found across brain areas and across species (Braganza & Beck, 2018). Circuit motifs have been suggested to represent the physical substrates of ‘computational primitives’ (*The Atoms of Neural Computation*, n.d.). There a are small number of fundamental, recurring circuit motifs (e.g., feed-forward, feedback, lateral inhibition, etc.) (Luo, n.d.). And so, a new conceptual approach in this paper is to ask how are specific circuit motifs assembled during development. In this study, we identified a new feed-forward circuit motif (*Figure 5*). Generally, feed-forward motifs are characterized as a pattern of connectivity in which one neuron (or neuron type) provides both direct and indirect input onto a second neuron (or neuron type). Feed-forward circuit motifs are common, found in many animals (e.g., nematodes, insects, mouse) and in many brain regions (e.g., somatosensory systems, olfactory systems, neocortex) (Schafer, 2016)(Anton & Homberg, 1999)(Harris & Shepherd, 2015). Thus, feed-forward motifs are fundamental to neural signal processing. Feed-forward motifs can be further subdivided into feed-forward inhibitory or feed-forward excitatory circuit motifs, depending on the transmitter types of the neurons involved (Luo, n.d.). Notably, anatomically the pattern of synaptic connectivity among neurons is the same in either motif subtype, and so in this study we do not distinguish between the two.

Here, we find that in general, early-born ELs get direct excitatory synapses from chordotonal sensory neurons and indirect (excitatory or inhibitory) input from chordotonals via Bains and Ladder interneurons (*Figure 5)*. Notably, there are differences in connectivity patterns among early born ELs. For example, although all early-born ELs get direct input from chordotonals and Ladders, A08j1-3s get left-right symmetrical inputs, whereas A08x and A08m get asymmetrical inputs *(Figure 5*). This could correspond to functional differences. For example, A08x and A08m may be involved in left-right asymmetrical signal detection. Teasing apart the diversity of computations performed by each individual EL interneurons is the domain of future studies.

Because we characterized the development of a feed-forward circuit in Drosophila we have information that allows us to search for similar circuits in other insect species. For example, in Drosophila, EL interneurons express the transcription factor, Even-skipped (Eve, *Figure 1, S1*). Eve is also expressed in ELs interneurons in and Locusts and other insects (Bevan & Burrows, 2003). Further, in Drosophila, the MNB lineage generates H-shaped, inhibitory Ladder interneurons, which get input from sound/vibration sensitive, chordotonal sensory neurons (*Figures 5E, H, Table S3, 8*). Similarly, in Locusts, the MNB lineage generates H-shaped, inhibitory interneurons, which encode sound stimuli (Thompson & Siegler, 1991). Further, in Drosophila and locusts, MNB interneuron progeny express the transcription factor Engrailed (Schmid et al., 1999)(Kearney et al., 2004)(Patel et al., n.d.). Thus, the neuronal components of the Ladder-to-early born EL circuit are conserved, which raises the possibility of circuit level conservation.

### Deepening our understanding of temporal cohorts

A major unanswered question in developmental neuroscience is how the mechanisms that generate neuronal diversity contribute to the formation of functional neuronal circuits (Sagner & Briscoe, 2019). Part of the answer lies in the observation that temporal cohorts are subunits of lineages, linked both to larval circuit anatomy/function as well as to the embryonic gene expression programs that generate neuronal diversity (Meng et al., 2019)(Meng et al., 2020)(Seroka & Doe, 2019). Thus far, temporal cohorts had been looked for and found in eight (of 30) lineages in the nerve cord (Wreden et al., 2017)(Meng et al., 2019)(Meng et al., 2020)(Mark et al., 2021). Here, we identify two additional temporal cohorts—Basins and Ladders—in two additional lineages—NB3-5 and MNB (*Figures 6-9*), bringing the number to ten. This underscores the idea that temporal cohorts are common.

A major unresolved question about temporal cohorts was to what extent are temporal cohort borders associated with sharp changes in connectivity. Previous studies had identified temporal cohorts and linked them with function and connectivity (Wreden et al., 2017)(Meng et al., 2019)(Meng et al., 2020)(Mark et al., 2021). However, these studies lacked the resolution to distinguish between a “graded” and “sharp” wiring transition models. We identified four temporal cohorts (early-born ELs, late-born ELs, Basins, Ladders) in three lineages (NB3-3, NB3-5, MNB) all of which have sharp changes in connectivity correlated with temporal cohort borders (*Figures 4,7-8*). For the Basin temporal cohort, we note that Basin have similar connectivity patterns with two additional NB3-5 progeny, one of which is likely to have birth order adjacent to Basins (*Figure 7G-H*). This suggests that the Basin temporal cohort contains neurons with morphologies and gene expression patterns distinct from Basin interneurons, underscoring the idea that temporal cohorts are subunits of hemilineages defined by birth within a tight time window rather than by neuronal morphology per se. Further, we note that within a temporal cohort, sequentially-born neurons are often, but not always, the most similar in terms of connectivity. For example, among the early-born EL temporal cohort, the fifth-born neuron (A08m) is more similar to the first-born neuron (A08x) than its temporal neighbor (A08j2), the fourth-born EL. Thus, our data, combined with previous studies show how temporal cohorts are developmental units related to circuits both at the functional and anatomical levels.

A second major unresolved question about temporal cohorts is the extent to which they are copies of each other. One reason this is an interesting question relates to circuit evolution. For the evolution of gene function, a popular model is a “duplicate and diverge” model. Similarly, temporal cohorts could be duplications within a lineage whose function could then diverge thereby driving circuit evolution. Such an idea motivated us to ask to what extent temporal cohort are copies. For example, do early-born ELs processes chordotonal stimuli in a manner identical to how late-born Els process proprioceptive stimuli? Our data suggest a more complex picture. Early-born and late-born ELs differ in their connectivity patterns—including left-right and following pair similarities (*Figure 4B*) and inter-and intra-segmental connectivity patterns (*Figure 9G-H*), suggesting that early-born and late-born ELs might differentially process information. Independent of the evolutionary implications, our data reveal previously unknown diversity in the structure of connectivity among neurons within adjacent temporal cohorts, which may indicate a diversity in underlying circuit assembly mechanisms.

### A feed-forward circuit is assembled from a select set of temporal cohorts

It has been hypothesized that nerve cord circuits are assembled by preferential connectivity between distinct temporal cohorts (Meng & Heckscher, 2021). Our data provide strong experimental support for this hypothesis. Specifically, we find that interneurons from three temporal cohorts wire together to form a feed-forward circuit—early-born ELs from NB3-3, mid-to-late born Basins from NB3-5, and late-born Ladders from MNB (*Figures 5-7*). Furthermore, other neurons from NB3-5 and MNB lineages do not synapse on early-born ELs.

Notably, one other study provided limited supported for the hypothesis that Drosophila nerve cord circuits are assembled by preferential connectivity between distinct temporal cohorts. This study focused on the Jaam-to-late-born EL-to-Saaghi circuit (Heckscher et al., 2015). Jaams are later-born interneurons in the NB5-2, Notch OFF hemilineage, and Saaghis are later-born interneurons in th NB5-2 Notch ON hemilineage (Mark et al., 2021). These data raised several possibilities: 1) There could be global alignment among lineages (e.g., all NB3-3 neurons get synapses from NB5-2 neurons). 2) Notch ON/OFF pairs of neurons might be pre-/post-synaptic partners of neurons within a temporal cohort. 3) Birth-order matched temporal cohorts might selectively wire together (e.g., early--early and late-late connectivity). However, our data demonstrate that none of these possibilities are generally true. Instead, we find a diversity in the manner in which temporal cohorts associate. The one consistent theme is that a limited number of temporal cohorts selectively interconnect.

It is of note that there are neurons that synapse onto early-born ELs, which we have not grouped into temporal cohorts. These include six neurons that synapse onto early-born ELs multiple (>10) times (Table S2). We cannot group these neurons into temporal cohorts because we do not yet have the tools to determine their lineage/birth order. And so, we cannot currently distinguish between the possibilities that these neurons are not in temporal cohorts versus the possibility that there exist more temporal cohorts that wire with the early-born ELs. By guessing lineage of origin based on anatomy alone, we favor the idea that at least some additional temporal cohorts synapse onto early-born ELs, but perhaps not as extensively as the Ladders and Basin temporal cohorts.

### Circuit inputs are born after circuit outputs

For most of this Discussion, we labeled neurons as “early-born” and “late-born”. These labels refer to the birth order of neurons within a lineage. However, these labels do not refer to the absolute time at which a neuron is born. This is because in the Drosophila nerve cord, neuroblasts are generated over a large span of embryogenesis (J. Broadus et al., 1995). Thus, early-born neurons from one lineage can be generated at the same time as later-born neurons from a different lineage. The absolute birth time of a neuron is related to the environmental context into which it is born (for example, what external signals or other neurons are present). Our study is unique in that it determined both neuronal birth order and birth time. From this analysis, we learn that early-born EL interneurons (i.e., circuit outputs) are born before Ladder interneurons and Basin interneurons (i.e., circuit inputs) (*Figure 10*). More generally, this suggests that for assembly of feed-forward circuit motifs, pre-synaptic interneurons are born after their post-synaptic partners. Further, it is possible that this principle extends to the assembly of other circuits that contain motor neurons. Motor neurons are always circuit outputs (to muscle) and are among the first neurons to be born during neurogenesis (Meng & Heckscher, 2021). Moreover, this pattern holds true in both Drosophila nerve cord and spinal cord (Schmid et al., 1999)(Fetcho & McLean, 2010). Therefore, our data raise the possibility that a fundamental rule for circuit assembly is that feed-forward circuits are assembled sequentially from circuit output to circuit input.

## Materials and Methods

### Drosophila strains and culture

Fly stocks were maintained at 25°C on standard cornmeal molasses medium. Experimental crosses were set up at the indicated temperatures. See below for details. See Table S4 for details of fly lines.

### Immunohistochemistry

Larvae at different stages were dissected in Baines’ solution. The dissected larval brains were adhered to poly-lysine coated coverslip and fixed for 7 min in 4% paraformaldehyde solution (Electron Microscopy Services, Hatfield, PA) as previously described (Heckscher et al., 2014). Larval brains were then washed 3 times with phosphate buffered saline containing 0.1% Triton X-100 (PBT), blocked 1hr in PBT containing 2% normal goat serum (NBT), and stained with primary antibody in PBT at 4°C overnight. After washing, samples were stained with secondary antibody at room temperature for 1hr, washed again, and applied with increasing percentatges of ethanol in water (30, 50, 70, 95, 100) series to replace PBT. Samples were then immersed into xylene for clearance, and then mounted in DPX (Sigma, MI). Images were acquired using either a Zeiss LSM 800 confocal microscope, and were processed and analyzed using Image J. For list of primary antisera see Table S3. Secondary antibodies were obtained from ImmunoResearch (Bar Harbor, ME) and were used at a 1:400 dilution.

### Generation of the *pPL[KD]* transgene

The KDR based permanent labeling plasmid, *pPL[KD]* (*nSyb* promoter and 5’UTR-KDRT-stop-KDRT-IVS-Syn21-GAL4-VP16-p10 3’UTR) was constructed in the backbone of nSybKOG (gift from Dr. Tzumin Lee) by restriction enzyme mediated molecular cloning (Awasaki et al., 2014). The GAL4 coding region in nSybKOG was replaced by IVS-Syn21-GAL4-VP16-p10 3’UTR sequence. The IVS and p10 fragments were derived from the vector pJFRC81 (Addgene #36432), Syn21 sequence (Pfeiffer et al., 2010) was added by PCR amplification, and GAL4-VP16 was derived from the vector pBPGAL4.2::VP16Uw (Addgene #26228). The resulting *pPL[KD]* plasmid was incorporated into the docking site VK27 by integrase-mediated site-specific integration, performed by GenetiVision Corporation (Houston, TX). Details of molecular cloning and construct sequence are available upon request.

### Generation of the EMS-zp-AD transgene

The pEntr-EMS plasmid was generated from a HiFi reaction (NEB) using the pEntr3c Gateway cloning (Invitrogen) plasmid digested with KpnI and NotI restriction endonucleases and 4 PCR reactions spanning the EMS promoter sequence found in Estacio-gomez et al (Estacio-Gómez et al., 2013). The primer pairs used were: 1) CGACTGGATCCGGTACCccagacagaactccatactccac and gtcgttaaacAAATGAATTGCCATAAGCG; 2) caattcatttGTTTAACGACCAACGCTC and tccggatggtCGAGCGGGATTTATGAGC; 3) atcccgctcgACCATCCGGATCTGGGCAAAAC and aatgaaaaccGTAAAAAATGCAGCCAACAAAGGG; 4) cattttttacGGTTTTCATTCCTTTTTGCG and GTCTAGATATCTCGAGTGCGGCCGCgtgtagtatggccgtcttctttgc. The pEntr-EMS was combined with the Gateway cloning destination vector pBPzpGal4AD using an LR reaction to generate pBP-EMSzpAD.

### **t**s-MARCM experiments

In this study we used ts-MARCM to determine both neuronal birth order and to estimate neuronal birth time. *<u>Birth order</u>* is determined in reference to other neurons within a given lineage, e.g., first-born, second-born. Birth order is also often referred to as a neuron’s temporal identity. Neuronal temporal identity is assessed by expression of temporal transcription factors. Consequently, determining birth order can be used to infer temporal transcription factor expression. *<u>Birth time</u>* refers to when during neurogenesis a neuron is born. Birth timing is usually reported in units of minutes or embryonic stage. Determining neuronal birth timing provides context into which a neuron is born (e.g., availability of other early-born neurons, transient signaling cues). In Drosophila embryos, neuron birth timing and birth order are not identical because during embryogenesis neuroblasts start to divide at different times. For example, NB3-5 is generated relatively early in embryogenesis and generates a large number of neurons, whereas MNB is generated much later and generates fewer neurons. And so, MNB can be generating first-born neurons at the same time that NB3-5 is generating much later-born neurons. We tried to birth date neurons using photo-activation and EdU/BrdU labeling, but were unsuccessful. So, instead, we estimated Basin birth timing using a series of carefully timed heat shocks to generate ts-MARCM clones. See below for details.

### EL-GAL4

We used the pan-EL driver, *EL-GAL4*, which is expressed in all ELs (Heckscher et al., 2015). Embryos of the genotype *hsFLP; UAS-rCD2.RFP, UAS-GFPi, FRT40A/ UAS-mCD8-GFP, UAS-rCD2i, FRT40A; EL-GAL4/ pPL[KD].* Addition of the pPL[KD] was required to amplify expression from *EL-GAL4 (Figure 1C, S1)*. This optimization step was required to drive high enough levels of RNAi constructs that are at the core of the ts-MARCM strategy (e.g., UAS-rCDi). High RNAi levels are needed such that “leaky” expression of membrane reporters (e.g., UAS-rCD3.RFP) are suppressed. Also, high levels of membrane-tethered reporter protein are needed in order to observe the detailed morphology of the entire neuron arbor.

*<u>Birth order (Figure 1C-E; S1).</u>* To determine the birth order of A1 ELs, embryos were collected for 2-4 hours intervals on apple juice plates. Collections were aged for 5 hours to 13 hours after egg collections. The collected samples were exposed to 37°C heat shock for 20 to 25 minutes. Heat shock stochastically induces FLP expression, which ultimately generates ts-MARCM clones. Heat shocked samples were incubated at 29 °C to boost GAL4 activity. Larvae were dissected at late L1 stage. Individual Al EL clones were visualized in samples with extremely sparse labeling (usually 1 to 2 clones per CNS) in order to obtain a clear morphology. For each EL clone, the generic EL identity of every labeled neuron was confirmed by Eve protein staining. Then the specific identity of each singly-labeled EL neuron was determined by matching the morphology to connectome data (see below). Singly-labeled neurons are the sole neuron expressing one fluorophore (either GFP or RFP). The remaining neurons expressed the opposite fluorophore, and we call these alternatively-labeled neurons. To determine birth order of singly-labeled ELs, we counted the number of alternatively-labeled Eve(+) neurons. As a confirmation of assigned birth order we also counted the number of unlabeled ELs (i.e., GFP[-], RFP[-], Eve[+]). To order A08e3, A08e2, and A083e1, we use the following logic. In our data set, A08e2 was always singly-labeled, and so A08e2 is mostly likely that last-born EL. Further, Figure 1C (and other examples) clearly shows both that A08e3 is the third from last-born and that later-born neurons are local. Therefore, we conclude A08e1 must be the second from last-born. The ts-MARCM clonal experiments were performed repeatedly until we assigned the birth order of all the A1 ELs. A total of 58 CNS were scored.

*<u>Birth time (Figure 10).</u>* The logic behind using ts-MARCM to determine birth timing is as follows: If a heat shock treatment occurred in embryos at the developmental stage either prior to during the cell division that generated a particular neuron type, then a clone will be induced. On the other hand, if the heat shock treatment occurred in embryos at the developmental stage after a neuroblast divides to give rise to a particular neuron, no clones will be generated. Therefore, determining birth timing requires more accurate staging of the developing embryos compared to experiments used to determine birth order.

Notably, in our ts-MARCM experiments there is a delay between when the heat shock is delivered to the embryo and the time when FLP recombinase protein has been produced and is active in dividing cells. To estimate this delay, we used ELs to “calibrate” our assay. Specifically, we crossed ts-MARCM transgenes to *EL-GAL4.* We let flies lay eggs for one hour, then aged the egg collection for 7hr or 9hr before delivering a heat shock. In embryos aged 7 hours, the most frequently produced single-labeled EL is A08m. A08m is the 5th-born EL. The 5th born EL is formed at ∼9 hours of development (Tsuji et al., 2008)(Gunnar et al., 2016)(Demilly et al., 2011). In embryos aged 9 hours, the most frequently produced single labeled EL was A08s. A08s1 is the 8th born EL. The 8th born EL is formed at ∼11 hours of development (Gunnar et al., 2016)(Tsuji et al., 2008). From these data, we infer that in our assay there is a ∼2 hour delay between the time of heat shock and the time of FLP-induced recombination in a dividing cell.

#### BASIN-GAL4 *(Figure 6)*

To determine the birth order and birth timing of Basins, we used the pan-Basin driver, *72F11-GAL4*. Embryos of genotype *hsFLP; UAS-rCD2.RFP, UAS-GFPi, FRT40A/ UAS-mCD8-GFP, UAS-rCD2i, FRT40A; 72F11-GAL4/+* were collected for 1 hour on apple juice plates. Collections were aged for 6 hours to 13 hours after egg collections. The collected samples were exposed to 37°C heat shock for 20, 30, or 35 minutes. Heat shocked samples were then incubated at 29°C to boost GAL4 activity. Larvae were dissected at late L1 stage. We scored a total of 113 clones.

*<u>Basins are lineage-related.</u>* We generated low levels of recombination at early times in development by providing 20-minute heat shock at 6 or 7 hrs after collection. A total of 12 CNSs were scored, and of those seven had one or two clones labeled. Of these seven CNSs, at least one of the two clones was a four-cell Basin clone.

*<u>Birth order.</u>* Individual Basin clones were visualized in samples with sparse labeling in order to obtain a clear morphology. For each Basin clone, we identified the singly labeled neuron by matching it to connectome data. We birth ordered the singly-labeled Basin by counting the number of alternatively-labeled neurons in the clone.

*<u>Birth timing.</u>* We determined when Basins were born. Basins are progeny of NB3-5. NB3-5 divides from stages 9 to 16 (Schmid et al., 1999). Each NB3-5 division gives rise to a ganglion mother cell, which divides once to produce two progeny (Monedero Cobeta et al., 2017) (Moris-Sanz et al., 2014). For embryos of each age, we scored the number of hemisegments displaying the following clonal morphologies:

- <u>0:4 clones</u>. In 0:4 clones, all Basins have the same label, indicating the heat shock occurred before NB3-5 divided to generate the first Basin. 0:4 was the most common clone type for heat shocks applied at 6 or 7 hrs (*Figure S4J*). This corresponds to stage 12 embryos.
- <u>1:3 clones</u>. In 1:3 clones, three Basins are labeled with one marker, and the other Basin is singly-labeled with the other marker. This indicates the heat shock occurred as NB3-5 was dividing to make the first Basin. <u>1:2 clones.</u> NB35 is dividing to make the second Basin. <u>1:1</u> <u>clones</u>. NB3-5 dividing to make third Basin. Clones of the 1:3, 1:2, and 1:1 types were most frequently observed in 8, 9 and 10 hr heat shock experiments. This corresponds to stages 13-15.
- <u>0:1 clones</u>. In 0:1 clones heat shock occurred either as NB3-5 was dividing to make the final Basin, or as a GMC was dividing to make a Basin. In embryos heat shocked at 11 hr we find only 0:1 clones, suggesting at this time point only GMCs are dividing to produce Basins. This corresponds to late stage 15. We note that there are examples of 0:1 clones in all experiments that provided heat shocks from 6 hr to 11hrs. This could indicate some stochastic labeling. However, we also note that in embryos heat shocked at ages 12 and 13 hours, Basin clones were never observed. Lack of clones indicates no cells were dividing to generate Basins.

Thus, peak Basin production occurs in embryos that have been heat shocked after they were aged 8 to 10 hours. Taken together with the idea that there is an estimated delay of two hours from time of heat shock to FLP-induced recombination (see EL-GAL4 section above), this suggests that, in general, the majority of Basin interneurons are generated from NB3-5 between 10 and 12 hours of development, or late stage 13 to stage 15. Furthermore, these data suggest that peak Basin production is likely to occur within a two-hour time window. This means in two hours NB3-5 produces four Basins, or one Basin every 30 minutes. In Drosophila larvae, neuroblasts are thought to divide with a frequency of once every 45 minutes. Although, we cannot definitively rule out the idea that production of one or more neuron type could sneak in, these data are consistent with the idea that Basins are continuously-born.

To gain confidence that our heat shock experiment timing was robust across experiments and genotypes, we performed a similar set of experiments with *Basin-1-GAL4* (aka, *78F07-GAL4*) (*Figure 10*). *Basin-1-GAL4* labels only one Basin, which is distinct from *Basin-GAL4*, which labels all four Basins. Our *Basin-1-GAL4* and *Basin-GAL4* datasets are in general agreement, suggesting that our assay is robust across different genotypes and experiments. See below for details.

#### BASIN-1-GAL4 and LADDER-D-GAL4 *Figure 10*

We used the neuron-specific drivers, *78F07-GAL4* and *20B01-GAL4*. These drivers expressed solely in Ladder D and Basin 1, respectively, but not in other neurons within the same lineage. Embryos of genotype *hsFLP; UAS-rCD2.RFP, UAS-GFPi, FRT40A/ UAS-mCD8-GFP, UAS-rCD2i, FRT40A; 78F07-GAL4* and *hsFLP; UAS-rCD2.RFP, UAS-GFPi, FRT40A/ UAS-mCD8-GFP, UAS-rCD2i, FRT40A; 20B01-GAL4*.

*<u>Birth timing.</u>* To determine the birth timing of Ladder D and Basin 1, embryos were collected within an hour interval on apple juice plates. They were aged for 6 hours to 13 hours after egg collection. The collected samples were exposed to 37°C heat shock for 30 minutes, and then incubated at 29 °C to boost GAL4 activity. Larvae were dissected between late second larval instar and early third larval instar, within which the expression of the GAL4 drivers reach the peak of their intensities. Both Ladder D and Basin 1 can be clearly identified by the morphology (*Figure 10C-D*). The segmental identity was distinguished by Eve protein staining. We scored neuronal clones in segments A1 to A7 based on the assumption that individual neurons should develop roughly at the same time in different abdominal segments, i.e. Basin 1 in A1 and in A7 should form roughly simultaneously. This broader scope of scoring, together with the stronger clonal induction maximized our capability to capture any larval CNS that had at least one stem cell or GMC underging division out of 7 abdominal segments (or 14 hemi-segments). The last time point that we could catch larvae CNS with a ts-MARCM clone was last time point at which there could be a division to generate Ladder D or Basin 1. A total of 57 CNSs were scored.

#### BASIN-GAL4 and EL-GAL4 *Figure 10*

We used both the pan-Basin driver, 72F11-GAL4, and the pan-EL driver, EL-GAL4. Embryos of genotype hsFLP; UAS-rCD2.RFP, UAS-GFPi, FRT40A/ UAS-mCD8-GFP, UAS-rCD2i, FRT40A; 72F11-GAL4/EL-GAL4

*<u>Birth timing.</u>* To determine the relative birth order of Basins and ELs, embryos were collected for 1 hour on apple juice plates. Collections were aged for 7 hr or 9 hr after egg collections. The collected samples were exposed to 37°C heat shock for 33 minutes. The analysis was identical to that described for *Basin-GAL4* above. 29 CNSs were scored.

### Multi-color FLP Out (*Figure 9*)

We labeled single neurons using Multi-Color FLP Out (MCFO) (Nern et al., 2015). MCFO stochastically labels with epitopes the membranes of cells within a GAL4 pattern. To obtain single cell clones, adult flies laid for 24 hours on apple juice caps. Caps were heat shocked in a water bath at 37 °C to 39°C for 15 to 30 minutes and incubated at 25°C for 4 to 5 hours. LI larvae were dissected. Their brains were stained for HA, Flag, and V5 epitopes to visualize single cell clones. Larvae were also stained for Eve protein to confirm the identity of each single cell clone as an EL interneuron, and to assign segmental identity to each clone. We generated >100 single cell clones and saw each neuronal morphology in a minimum of two separate larvae. Each clone was analyzed in dorsal and posterior views

### Identification of stem cell lineage of Basins and Ladder D *(Figure 7B-D, 8C-E)*

To assign neurons to a specific neuroblast lineage, larvae of genotype *Basin-1-LexA/UAS-mCherry.NLS; NB3-5-GAL4/ LexAop2-CsChrimson-mVenus* were used. Where *NB3-5-GAL4* was one of three different GAL4 lines (*49C03-GAL4, ham-GAL4*, *59E09-GAL4*). We used three different GAL4s to label NB3-5 because any one GAL4 line might have un-expected off target labeling. We reasoned that each line should have different off target labeling, and so if we saw co-localization of markers in all three cases, this could be taken as strong evidence that Basin 1 was in the NB3-5 lineage. Lines also included FLP based permeant labeling (UAS-FLP, *actin* promoter and FRT-stop-FRT-GAL4) which labels all progeny from a given neuroblast. Larvae were raised at 29 °C on the apple juice plate. The larvae were then dissected in the L3 stage for further staining and imaging processing.

We used a similar logic to determine neuroblast origin for Ladders. Larvae of genotype *Ladder-D-LexA/UAS-mCherry.NLS; MNB-GAL4/ LexAop2-CsChrimson-mVenus* were used. Where *MNB-GAL4* was one of three different GAL4 lines (*13G03-GAL4, 70D06-GAL4*, and *65H11-GAL4*).

### Calcium imaging *(Figure 5J-L)*

For calcium imaging experiments, all larvae were within 6 hr of age on the day of recording and collected 48 to 54 hr after hatching. Larvae expressing GCaMP6m were rinsed with water and placed ventral side up on agarose pads with a 22 mm x 22 mm coverslip placed on top. Pads were made by pouring 3% agarose into a well. Recordings began with a 30 second period of no stimulus followed by a 30 second period of sound stimulus and ending with a final 30 second period of no stimulus. A Visaton FR12, 4 Ohm speaker (5 inches diameter) and a PYLE PCA2 stereo power amplifier was used to project sound. For further details, refer to Marshall and Heckscher (Marshall & Heckscher, 2022). Images were acquired on a Zeiss LSM 800 confocal microscope using 0.1%–0.2% 488 nm laser power with the pinhole entirely open. Images were acquired at 3 frames per second using a 10X (0.3 NA) or 20X (0.8 NA) objective. The calcium signal was continuously collected before, during, and after the stimulus. Extracting changes in GCaMP6m fluorescence amplitude was done using Fiji as in Marshall and Heckscher (Marshall & Heckscher, 2022). A region of interest (ROI) that included the larval nerve cord was manually drawn, and the mean fluorescence within the ROI was calculated for each time point.

### Connectome analyses

The connectome dataset used in this study is a CNS reconstruction from a 6 hr old first instar larva, which was described by Ohyama et al., 2015.

#### Identifying NB3-3A1L/R neurons in the connectome (Figures 1, S1, 2, Table S1)

In the connectome, the following ELs had been previously identified (A08x, A08m, A08c, A08s, A08e1, A08e2, A08e3)(Wreden et al., 2017)(Heckscher et al., 2015). To identify other neurons in the lineage, we used the following logic: NB3-3A lineage has thirteen neurons, including two non-EL neurons, one of which is a motor neuron and eleven EL interneurons (Tsuji et al., 2008)(Schmid et al., 1999). We found that all EL interneurons form a bundle (*Figure 2B-C*), and that the two non-EL neurons are also part of that bundle (*Figure 2A*). Therefore, in the connectome, we looked for a bundle that contained thirteen neurons and that included all previously-identified EL interneurons. This bundle is shown in Figure 2C. It contains two non-EL neurons-a motor neuron, an undifferentiated neuron, all previously identified ELs, and several interneurons that were candidates to be ELs. We confirmed these candidate neurons were indeed ELs by making single neuron clones with *EL-GAL4* driving ts-MARCM constructs and co-staining clones with anti-Eve (*Figure 1C, S1*). For details on matching neurons in light level and connectome level datasets see Heckscher et al., and Wreden et al. (Wreden et al., 2017)(Heckscher et al., 2015). This confirmed the identity of A08j1, A08j3, A08j2, and A08o as ELs. While this manuscript was in preparation Mark et al., partially characterized multiple lineages, including NB3-3 (Mark et al., 2021). In agreement with our assignments, Mark et al suggest that A08j1-A08j3 and A08o are ELs. Notably, our naming scheme for ELs differs slightly from that used by Mark et al.. In our naming scheme, we use the first published name for each EL. Details can be found in Table S5.

#### Identifying NB3-5A1 neurons in the connectome (Table S3, Figure 7E)

In the connectome, Basin 1-4 in segment A1 had been previously identified (Ohyama et al., 2015a)(Jovanic et al., 2016). To identify other neurons in the lineage, we used the following logic: First, we found all neurons that bundled with Basin neurons (total of 34). Of these, 10 neurons had a medial trajectory upon entering the neuropil. We excluded these neurons from further analysis for two reasons. 1) They are of a number and morphology that is consistent with their being from NB2-4 (Schmid et al., 1999). 2) The NB3-5 lineage is reported to have the capability of generating as many as 36 neurons (Monedero Cobeta et al., 2017). But, by late embryonic stage 17, only 19-24 are still found in abdominal segments (Schmidt et al., 1997). Excluding the 10 putative NB2-4 neurons from the lineage bundle, left 24 neurons, which matched the number reported by Schmidt. Of these neurons, 5 were undifferentiated—with neurites ending prematurely with no synapse input or output. We excluded these from further analysis. The remaining 19 neurons fell into to broad morphological categories—neurons with dorsally-projecting neurites and neurons with ventrally-projecting neurites. Recently, for many lineages in the Drosophila nerve cord, dorsally-projecting neurons were shown to belong to a Notch ON hemilineage and ventrally-projecting neurons to a Notch OFF hemilineage (Mark et al., 2021). NB3-5 produces two hemilineages (Monedero Cobeta et al., 2017) (Moris-Sanz et al., 2014). Of the remaining neurons in the lineage bundle, six “Drunken” neurons had neurite trajectories that diverge dorsally after entering the neuropil. We consider these to be potentially Notch ON neurons from the NB3-5 lineage and excluded them from our analysis. We were left with 13 neurons in the bundle, which also contained Basins. All of these neurons had ventral trajectory in the neuropil *(Figure 5C*). These 13 neurons were used for analysis (*Figure 7F-H, Table S3*).

#### Identifying Ladder neurons in the connectome (Table S2, Figure S5)

In the connectome, Ladder A-F in segment A1 had been previously identified (Jovanic et al., 2016). See Table S2 for Ladder inputs.

#### Neuronal reconstruction, finding synaptic partners, proof reading annotations

Once all the NB-3-3A1 neurons were identified, their skeletons were reviewed to greater than 90% with mainly areas in the brain unreviewed. Every skeleton with an input synapse onto a NB3-3A1L/R neuron was reconstructed in an attempt to generated complete neurons with cell bodies.

Upstream left/right neuron pairs were identified by mirror image morphology and synapse similarity. Upstream pairs that had input onto NB3-3A1L/R pairs at a greater than 4 or more synapses on one of the pair and 2 or more synapses on the other neuron of the pair were reviewed to greater than 80% (Table S1).

#### Calculating neuron birth order versus proximal neurite length (Figure 2E, 7F)

The distance was calculated using the CATMAID function “Measure the distance between two nodes.” The two nodes used were the cell body and a node chosen by eye where the skeleton entered the neuropil.

#### Calculating inputs onto NB3-3A1L/R by type (Figure 3A, Tables S1)

Input neurons onto NB3-3A1L/R neurons were broadly categorized into four types. Nerve cord interneurons had a cell body in segments T1-A10. Sensory neurons had no cell body, but instead had axons entering the nerve cord in bundles from the periphery. They also show a unique, electron-dense cytoplasm. Central brain and SEZ neurons had cell bodies anterior to the nerve cord. Unknown included fragments of neurons that could not be traced back to cell bodies.

#### Calculating sensory neuron inputs onto NB3-3A1L/R (Figure 3B)

A sensory neuron was determined to be chordotonal, proprioceptive, or other, based on morphology of the central axon (Grueber et al., 2007)(Heckscher et al., 2015)(Ohyama et al., 2015b).

#### Calculating major interneuron inputs onto NB3-3A1L/R (Figure 3C)

We displayed only “major” interneurons. We consider major interneurons to include the subset of nerve cord interneurons that matched the following criteria. 1) For hemisegemental homologs (i.e., left-right pairs of neurons), one neuron formed >3 synapses onto a NB3-3A1L or NB3-3A1R neuron and the other neuron formed >1 synapse onto the hemisegmentally homologous NB3-3A1L or NB3-3A1R neuron. 2) For unpaired midline neurons, the neuron formed >3 synapses with a NB3- 3A1L or NB3-3A1R neuron and formed >1 synapse with the hemisegmentally homologous NB3-3A1L or NB3-3A1R neuron. These neurons are listed as 4-2 in Table S1. In total, this was 174 nerve cord interneurons that synapses onto NB3-3A1L neurons, and 182 that synapse onto NB3-3A1R neurons.

#### Calculating synapses from nerve cord neurons onto early-born ELs (Figure 5A, G-I, Table S2)

Starting with all inputs to NB3-3A1L/A1R we found the subset of inputs onto early-born ELs (548 neurons, 2618 synapses). Of those, we identified interneurons and sensory neurons (see calculating inputs onto NN3-3A1L/R by type) (334 neurons, 2061 synapses). Of these neurons were binned into classes (chordotonals, Ladders, Basins, early-bornn ELs) based on previously-described morphological criteria (Ohyama et al., 2015b)(Jovanic et al., 2016).

### Distance analyses

#### Non-binary analysis (Figures 4A, C-E, 7G-H, 8F)

We treated inputs to a neuron as a vector, which contained the number of synaptic contacts. The order of inputs was the same for every pair of neurons. We then calculated the Euclidean distance in between the two inputs vectors. These distance vectors summarize how similar the two neurons’ inputs are. For the distance to be zero, not only would the input partners need to completely overlap, but the number of synapses contributed by each of these partners would need to be identical.

To test for significance, we created 100 surrogate vectors. We permuted the inputs of each neuron independently, preserving the number of inputs (in degree) but shuffling their identity. We then calculated the Euclidean distances of pairs in the shuffled vectors, building a distribution of Euclidean distances expected by chance. We normalized real data to shuffled data as follows: real-(mean[shuffled])/(standard deviation [shuffled]), which generated z-scores. Z-scores of greater than +1.96 or less than -1.96 were considered significantly more similar or different, respectively, than would be observed by chance (at the alpha = 0.05 level). Significant pairwise differences and similarities were defined as significant positive and negative (respectively) z scores of the real Euclidean distance, at p<0.05 (two-tailed).

#### Binary analysis (Figure 4B)

Since we observed a broad variation in synaptic strengths and input onto ELs (*Table S1*), to disambiguate the identity of inputs from their respective strength, we repeated distance analyses using input identity only. To do so, we treated inputs to a neuron as a binary vector, with 1 for the presence and 0 for the absence of input, respectively. Otherwise, analysis was identical to non-binary. Broadly speaking, results from binary and non- binary analyzes were similar. One notable exception is that binary analysis did not identify statistically significant differences (*e.g., Figure 4C*).

#### Left-right and following pair analysis (Figure 3B, 5J)

We examined the normalized (z) Euclidean distance scores obtained from binary connectomes regardless of their significance in two types of neuron pairs separately: Left-right pairs are comprised of the same neurons in the left and right hemisegment, for example, A08j2 A1L and A08j2 A1R. Following pairs are comprised of neurons that are born consecutively in the same hemisegment, for example, A08j3 A1L and A08j2 A1L. In *Figure 4B,* we pooled the scores of the two pair types over early-born, late-born and undifferentiated and motor neurons separately.

## Acknowledgements

We would like to thank the Janelia Fly EM Project Team for the gift of the EM volume, and the Janelia visitor program for their support. We would like to thank the following for contributing to EM annotation: Akria Fushiki, Albert Cardona, Andreas Schoofs, Antia Burgos, Aref Arzan Zarin, Avinash Khandelwal, Brittany Kemp, Casey M Schneider-Mizell, Chris Q Doe, Elizabeth Barsotti, Ingrid Andrade, Jamie Macleod, Julia Meng, Larisa Maier, Laura Herren, Maarten F Zwart, Xinyu Tang. Funding: NIH grants NS105748 (to ESH), EY022338 (to JNM), and T32 HD044164 (to ZDM)

---

**Supplemental Table 1. Inputs onto NB3-3A1L/R neurons**

With following tabs:

All inputs on NB3-3A1L/R neurons (Figure 3A-B) Cable length (Figure 2E)

Major interneuron inputs (Figure 3C)

NB3-3 to NB3-3 connectivity (Figure 9G-H)

**Supplemental Table 2. Inputs onto early-born ELs**

With following tabs: Summary (Figure 5B)

Inputs onto early-born ELs (Figure 5B) Input onto each early-born EL (Figure 5G-I)

All sensory neuron and interneuron inputs onto early-born ELs (Figure 5B) Lineage guesses (Figure 5B)

**Supplemental Table 3. Inputs onto Basins and Ladders**

With following tabs:

Basin A1 connectivity (Figure 7G-H) Basin names (Figure 7)

Ladder A1 connectivity (Figure 8F)

**Supplemental Table 4. Resources**

With the following tabs:

Genotypes used in this study

Fly lines

Antibody list

**Supplemental Table 5. Names for ELs in A1**

**Supplemental Figure 1.**
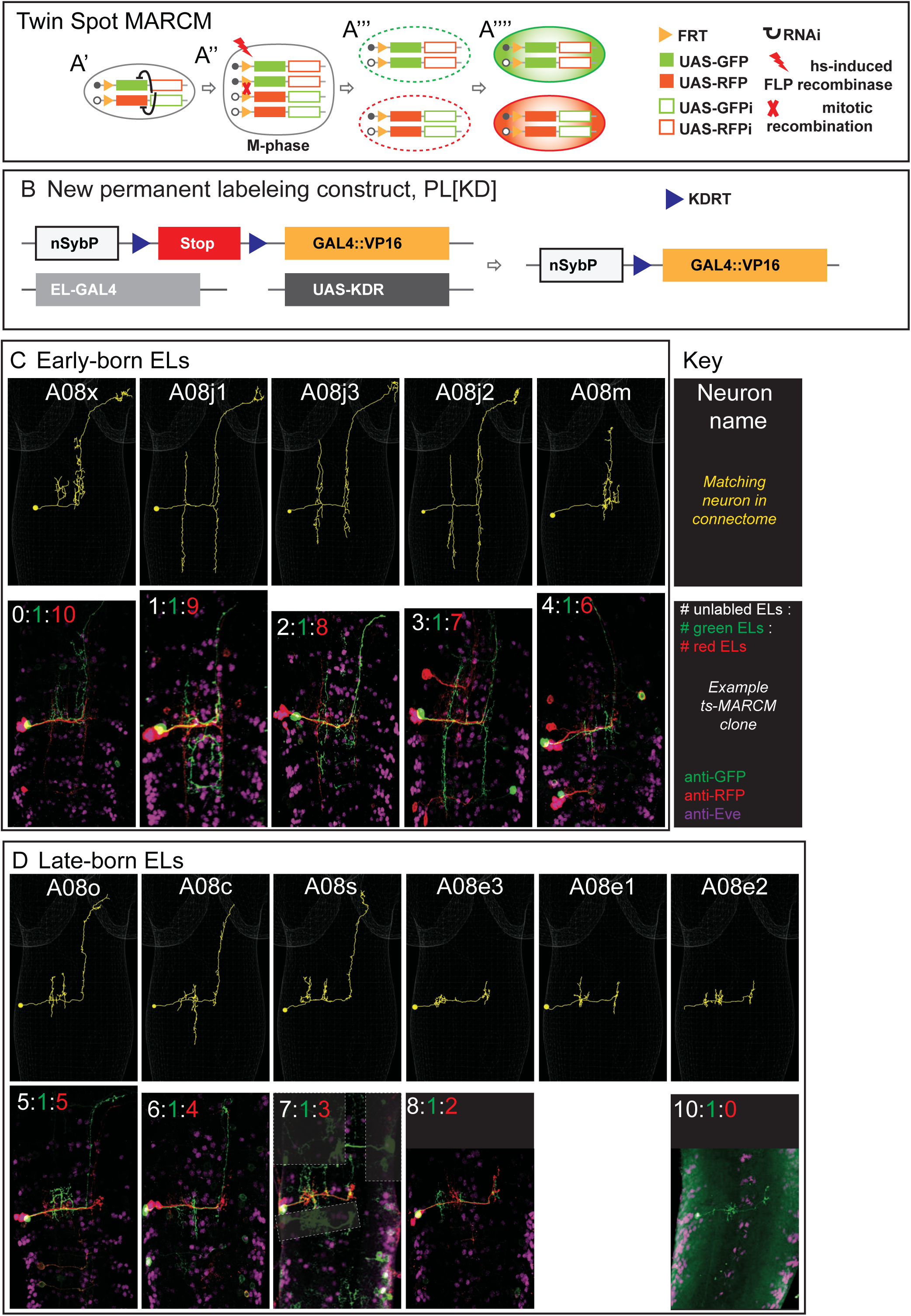
Twin Spot MARCM data. **A-B. Illustration of ts-MARCM** Our updated version of ts-MARCM system has four components. 1) It uses a pair of genetically modified chromosomes. On one chromosome is an FRT recombinase site (yellow triangle) followed by a *UAS-GFP* (solid green box) and a *UAS-RFP-RNAi* (hollow red box) construct. On the other chromosome is an FRT site followed by a *UAS-RFP* (solid red box) and a *UAS-GFP-RNAi* (hollow green box) construct. When cells are heterozygous for these chromosomes, the GFP- and RFP- RNAi constructs ensure repression of GFP and RFP protein expression, respectively (black curves, A’). 2) It has a heat-shock inducible FLP recombinase (red lightning bolt). By varying the heat shock protocol, we control both the timing and amount of FLP supplied. Heat shocks induce FRT-based chromosomal recombination in dividing cells (red X, M-phase cell, A’’). A subset of recombination events generate a pair of post-mitotic progeny, one of which is homozygous for the *UAS-GFP, UAS-RFP-RNAi* construct, and the other homozygous for the *UAS-RFP, UAS-GFP-RNAi* construct. In these cells, RNAi is no longer able to repress GFP or RFP expression (A’’’). 3) A cell-type specific GAL4 line, (e.g., *EL-GAL4,* light gray box in B) is used to drive expression of *UAS-RFP* or *UAS-GFP* (A’’’’). 4) To get robust ts-MARCM labeling in early stage larvae, it was often necessary to amplify GAL4 expression. To do so, we generated a new permanent labeling construct (B). Specifically, a neuron-specific nSyb promoter (white box) is upstream of a Stop (red box) flanked by KDRT (blue triangles) recombination sights. When the KDR recombinase (from UAS-KD, dark gay box) is supplied, the Stop is removed, and nSyb drives expression of a the new GAL4 (yellow box). This new GAL4 is the GAL4 DNA binding domain tethered to the strong transcriptional activator VP16. **C-D. Images of individually labeled ts-MARCM clones shown in birth order.** Early-born ELs are show in C, and late-born ELs are shown in D. An image key is shown to the right of C. Briefly, all images are shown in a dorsal view with anterior to the top. The neuron name is at the top of each image pair along with an example of the neuron in the connectome (yellow). The bottom of each image pair is an example of a clone stained with anti-GFP, anti- RFP, and anti-Eve. At the top of the example clone image is the number of unlabeled ELs (white), the number of ELs labeled in green, and the number of ELs labeled in red. Sometimes clones in other segments are lightly boxed over for clarity. For genotype see Table S4.

**Supplemental Figure 2.**
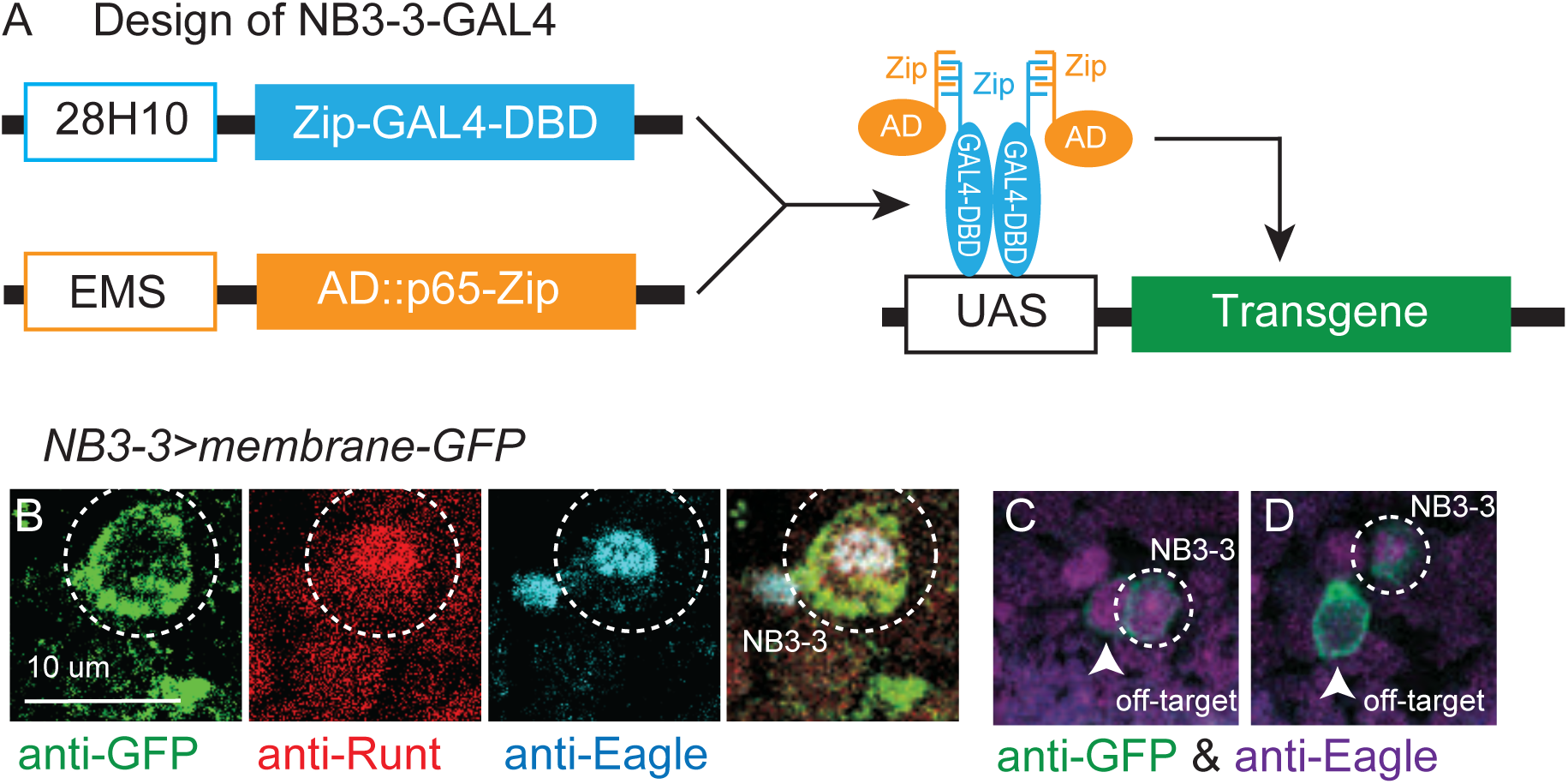
NB3-3-GAL4 line. **A. Illustration of NB3-3-GAL4 line** 28H10 (blue outlined box) and EMS (orange outlined box) are promoters that drive in different subsets of neuroblasts, whose expression overlaps specifically in NB3-3. Zip-GAL4-DBD encodes the DNA binding domain of the yeast transcription factor GAL4 (blue box). It binds the yeast UAS promoter. AD-Zip encodes an activation domain peptide that recruits transcriptional machinery to a promoter (orange box). When Zip-GAL4-DBD and AD-Zip are expressed in the same cell they activate expression of any transgene downstream of UAS promoter (green box). See methods for construction of EMS-zp-AD. **B-C. Images of NB3-3-GAL4 expression** B. *NB3-3-GAL4* labels NB3-3, which is selectively labeled by the overlap in expression of Runt (red) and Eagle (cyan). Dashed circle shows NB3-3. C. *NB3-3-GAL4* expresses in NB3-3 as well as a few, variable other neuroblasts. Eagle (magenta) labels four neuroblasts (NB3-3, NB6-4, NB2-4, NB7-3). Arrows point to NB-3 “off target” cells, dashed circles show a bone fide NB3-3. Single optical slices of stage 11 embryos are shown in dorsal view with anterior up NB3-3- GAL4 (dashed circle) drives membrane GFP. For genotype see Table S4.

## References

Anton, S., & Homberg, U. (1999). Antennal Lobe Structure. In B. S. Hansson (Ed.), Insect Olfaction (pp. 97–124). Springer. https://doi.org/10.1007/978-3-662-07911-9_5

Awasaki, T., Kao, C.-F., Lee, Y.-J., Yang, C.-P., Huang, Y., Pfeiffer, B. D., Luan, H., Jing, X., Huang, Y.-F., He, Y., Schroeder, M. D., Kuzin, A., Brody, T., Zugates, C. T., Odenwald, W. F., & Lee, T. (2014). Making Drosophila lineage–restricted drivers via patterned recombination in neuroblasts. Nature Neuroscience, 17(4), 631–637. https://doi.org/10.1038/nn.3654

Babski, H., Jovanic, T., Surel, C., Yoshikawa, S., Zwart, M. F., Valmier, J., Thomas, J. B., Enriquez, J., Carroll, P., & Garcès, A. (2019). A GABAergic Maf-expressing interneuron subset regulates the speed of locomotion in Drosophila. Nature Communications, 10(1), 4796. https://doi.org/10.1038/s41467-019-12693-6

Baumgardt, M., Karlsson, D., Salmani, B. Y., Bivik, C., MacDonald, R. B., Gunnar, E., & Thor, S. (2014). Global Programmed Switch in Neural Daughter Cell Proliferation Mode Triggered by a Temporal Gene Cascade. Developmental Cell, 30(2), 192–208. https://doi.org/10.1016/j.devcel.2014.06.021

Bevan, S., & Burrows, M. (2003). Localisation of Even-skipped in the mature CNS of the locust, Schistocerca gregaria. Cell and Tissue Research, 313(2), 237–244. https://doi.org/10.1007/s00441-003-0719-z

Braganza, O., & Beck, H. (2018). The Circuit Motif as a Conceptual Tool for Multilevel Neuroscience. Trends in Neurosciences, 41(3), 128–136. https://doi.org/10.1016/j.tins.2018.01.002

Broadus, J., Skeath, J. B., Spana, E. P., Bossing, T., Technau, G., & Doe, C. Q. (1995). New neuroblast markers and the origin of the aCC/pCC neurons in the Drosophila central nervous system. Mechanisms of Development, 53(3), 393–402.

Catela, C., & Kratsios, P. (2021). Transcriptional mechanisms of motor neuron development in vertebrates and invertebrates. Developmental Biology, 475, 193–204. https://doi.org/10.1016/j.ydbio.2019.08.022

Demilly, A., Simionato, E., Ohayon, D., Kerner, P., Garces, A., & Vervoort, M. (2011). Coe Genes Are Expressed in Differentiating Neurons in the Central Nervous System of Protostomes. PLoS ONE, 6(6), e21213. https://doi.org/10.1371/journal.pone.0021213

Doe, C. Q. (2017). Temporal Patterning in the DrosophilaCNS. Annual Review of Cell and Developmental Biology, 33(1), 219–240. https://doi.org/10.1146/annurev-cellbio-111315-125210

Doe, J. B. S. and C. Q. (1998). Sanpodo and Notch act in opposition to Numb to distinguish sibling neuron fates in the Drosophila CNS. 1–9.

Estacio-Gómez, A., Moris-Sanz, M., Schäfer, A.-K., Perea, D., Herrero, P., & Diaz-Benjumea, F.J. (2013). Bithorax-complex genes sculpt the pattern of leucokinergic neurons in the Drosophila central nervous system. Development, 140(10), 2139–2148. https://doi.org/10.1242/dev.090423

Fetcho, J. R., & McLean, D. L. (2010). Some principles of organization of spinal neurons underlying locomotion in zebrafish and their implications. Annals of the New York Academy of Sciences, 1198(1), 94–104. https://doi.org/10.1111/j.1749-6632.2010.05539.x

Grueber, W. B., Ye, B., Yang, C.-H., Younger, S., Borden, K., Jan, L. Y., & Jan, Y. N. (2007). Projections of Drosophila multidendritic neurons in the central nervous system: Links with peripheral dendrite morphology. Development, 134(1), 55–64. https://doi.org/10.1242/dev.02666

Gunnar, E., Bivik, C., Starkenberg, A., & Thor, S. (2016). Sequoiacontrols the type I>0 daughter proliferation switch in the developing Drosophilanervous system. Development, 143(20), 3774–3784. https://doi.org/10.1242/dev.139998

Harris, K. D., & Shepherd, G. M. G. (2015). The neocortical circuit: Themes and variations. Nature Neuroscience, 18(2), 170–181. https://doi.org/10.1038/nn.3917

Heckscher, E. S., Long, F., Layden, M. J., Chuang, C.-H., Manning, L., Richart, J., Pearson, J. C., Crews, S. T., Peng, H., Myers, E., & Doe, C. Q. (2014). Atlas-builder software and the eNeuro atlas: Resources for developmental biology and neuroscience. Development, 141(12), 2524–2532. https://doi.org/10.1242/dev.108720

Heckscher, E. S., Zarin, A. A., Faumont, S., Clark, M. Q., Manning, L., Fushiki, A., Schneider-Mizell, C. M., Fetter, R. D., Truman, J. W., Zwart, M. F., Landgraf, M., Cardona, A., Lockery, S. R., & Doe, C. Q. (2015). Even-Skipped+ Interneurons Are Core Components of a Sensorimotor Circuit that Maintains Left-Right Symmetric Muscle Contraction Amplitude. Neuron, 88(2), 314–329.

Isshiki, T., Pearson, B., Holbrook, S., & Doe, C. Q. (2001). Drosophila neuroblasts sequentially express transcription factors which specify the temporal identity of their neuronal progeny. Cell, 106(4), 511–521.

Jovanic, T., Schneider-Mizell, C. M., Shao, M., Masson, J.-B., Denisov, G., Fetter, R. D., Mensh, B. D., Truman, J. W., Cardona, A., & Zlatic, M. (2016). Competitive Disinhibition Mediates Behavioral Choice and Sequences in Drosophila. Cell, 167(3), 858–858.e19. https://doi.org/10.1016/j.cell.2016.09.009

Kearney, J. B., Wheeler, S. R., Estes, P., Parente, B., & Crews, S. T. (2004). Gene expression profiling of the developing Drosophila CNS midline cells. Developmental Biology, 275(2), 473–492. https://doi.org/10.1016/j.ydbio.2004.08.047

Li, H., Shuster, S. A., Li, J., & Luo, L. (2018). Linking neuronal lineage and wiring specificity. Neural Development, 13(1), 5. https://doi.org/10.1186/s13064-018-0102-0

Luo, L. (n.d.). Architectures of neuronal circuits. Science, 373(6559), eabg7285. https://doi.org/10.1126/science.abg7285

Mark, B., Lai, S.-L., Zarin, A. A., Manning, L., Pollington, H. Q., Litwin-Kumar, A., Cardona, A., Truman, J. W., & Doe, C. Q. (2021). A developmental framework linking neurogenesis and circuit formation in the Drosophila CNS. ELife, 10, e67510. https://doi.org/10.7554/eLife.67510

Marshall, Z. D., & Heckscher, E. S. (2022). The Role of Even-Skipped in Drosophila Larval Somatosensory Circuit Assembly. ENeuro, 9(1), ENEURO.0403-21.2021. https://doi.org/10.1523/ENEURO.0403-21.2021

Meng, J. L., & Heckscher, E. S. (2021). Development of motor circuits: From neuronal stem cells and neuronal diversity to motor circuit assembly. Current Topics in Developmental Biology, 142, 409–442. https://doi.org/10.1016/bs.ctdb.2020.11.010

Meng, J. L., Marshall, Z. D., Lobb-Rabe, M., & Heckscher, E. S. (2019). How prolonged expression of Hunchback, a temporal transcription factor, re-wires locomotor circuits. ELife, 8, 505. https://doi.org/10.7554/eLife.46089

Meng, J. L., Wang, Y., Carrillo, R. A., & Heckscher, E. S. (2020). Temporal transcription factors determine circuit membership by permanently altering motor neuron-to-muscle synaptic partnerships. ELife, 9, e56898. https://doi.org/10.7554/eLife.56898

Monedero Cobeta, I., Salmani, B. Y., & Thor, S. (2017). Anterior-Posterior Gradient in Neural Stem and Daughter Cell Proliferation Governed by Spatial and Temporal Hox Control. Current Biology, 27(8), 1161–1172. https://doi.org/10.1016/j.cub.2017.03.023

Moris-Sanz, M., Estacio-Gómez, A., Alvarez-Rivero, J., & Díaz-Benjumea, F. J. (2014). Specification of neuronal subtypes by different levels of Hunchback. Development (Cambridge, England), 141(22), 4366–4374. https://doi.org/10.1242/dev.113381

Nern, A., Pfeiffer, B. D., & Rubin, G. M. (2015). Optimized tools for multicolor stochastic labeling reveal diverse stereotyped cell arrangements in the fly visual system. Proceedings of the National Academy of Sciences of the United States of America, 112(22), E2967–76. https://doi.org/10.1073/pnas.1506763112

Ohyama, T., Jovanic, T., Denisov, G., Dang, T. C., Hoffmann, D., Kerr, R. A., & Zlatic, M. (2013). High-Throughput Analysis of Stimulus-Evoked Behaviors in Drosophila Larva Reveals Multiple Modality-Specific Escape Strategies. PLoS ONE, 8(8), e71706. https://doi.org/10.1371/journal.pone.0071706.s011

Ohyama, T., Schneider-Mizell, C. M., Fetter, R. D., Aleman, J. V., Franconville, R., Rivera-Alba, M., Mensh, B. D., Branson, K. M., Simpson, J. H., Truman, J. W., Cardona, A., & Zlatic, M. (2015a). A multilevel multimodal circuit enhances action selection in Drosophila. Nature, 520(7549), 633–639. https://doi.org/10.1038/nature14297

Patel, N. H., Martin-Blanco, E., Coleman, K. G., Poole, S. J., Ellis, M. C., Kornberg, B., & Goodman, C. S. (n.d.). Expression of engrailed Proteins in Arthropods, Annelids, and Chordates. 14.

Pearson, B. J., & Doe, C. Q. (2003). Regulation of neuroblast competence in Drosophila. Nature, 425(6958), 624–628. https://doi.org/10.1038/nature01910

Pfeiffer, B. D., Ngo, T.-T. B., Hibbard, K. L., Murphy, C., Jenett, A., Truman, J. W., & Rubin, G. M. (2010). Refinement of Tools for Targeted Gene Expression in Drosophila. Genetics, 186(2), 735–755. https://doi.org/10.1534/genetics.110.119917

Pinto-Teixeira, F., Koo, C., Rossi, A. M., Neriec, N., Bertet, C., Li, X., Del-Valle-Rodriguez, A., & Desplan, C. (2018). Development of Concurrent Retinotopic Maps in the Fly Motion Detection Circuit. Cell, 173(2), 485–498.e11. https://doi.org/10.1016/j.cell.2018.02.053

Sagner, A., & Briscoe, J. (2019). Establishing neuronal diversity in the spinal cord: A time and a place. Development, 146(22), dev182154. https://doi.org/10.1242/dev.182154

Schafer, W. (2016). Nematode nervous systems. Current Biology, 26(20), R955–R959. https://doi.org/10.1016/j.cub.2016.07.044

Schmid, A., Chiba, A., & Doe, C. Q. (1999). Clonal analysis of Drosophila embryonic neuroblasts: Neural cell types, axon projections and muscle targets. Development, 126(21), 4653–4689.

Schmidt, H., Rickert, C., Bossing, T., Vef, O., Urban, J., & Technau, G. M. (1997). The embryonic central nervous system lineages of Drosophila melanogaster. II. Neuroblast lineages derived from the dorsal part of the neuroectoderm. Developmental Biology, 189(2), 186–204.

Seroka, A. Q., & Doe, C. Q. (2019). The Hunchback temporal transcription factor determines motor neuron axon and dendrite targeting in Drosophila. Development, 146(7), dev175570. https://doi.org/10.1242/dev.175570

*The atoms of neural computation*. (n.d.). Retrieved March 31, 2022, from https://www.science.org/doi/10.1126/science.1261661

Thompson, K. J., & Siegler, M. V. S. (1991). Anatomy and physiology of spiking local and intersegmental interneurons in the median neuroblast lineage of the grasshopper. Journal of Comparative Neurology, 305(4), 659–675. https://doi.org/10.1002/cne.903050409

Truman, J. W., Moats, W., Altman, J., Marin, E. C., & Williams, D. W. (2010). Role of Notch signaling in establishing the hemilineages of secondary neurons in Drosophila melanogaster. Development, 137(1), 53–61. https://doi.org/10.7554/eLife.04493

Tsuji, T., Hasegawa, E., & Isshiki, T. (2008). Neuroblast entry into quiescence is regulated intrinsically by the combined action of spatial Hox proteins and temporal identity factors. Development, 135(23), 3859–3869. https://doi.org/10.1242/dev.025189

Wheeler, S. R., Stagg, S. B., & Crews, S. T. (2009). MidExDB: a database of Drosophila CNS midline cell gene expression. BMC Developmental Biology, 9(1), 56. https://doi.org/10.1186/1471-213X-9-56

Wreden, C. C., Meng, J. L., Feng, W., Chi, W., Marshall, Z. D., & Heckscher, E. S. (2017). Temporal Cohorts of Lineage-Related Neurons Perform Analogous Functions in Distinct Sensorimotor Circuits. Current Biology : CB, 27(10), 1521–1528.e4. https://doi.org/10.1016/j.cub.2017.04.024

Xu, H.-T., Han, Z., Gao, P., He, S., Li, Z., Shi, W., Kodish, O., Shao, W., Brown, K. N., Huang, K., & Shi, S.-H. (2014). Distinct lineage-dependent structural and functional organization of the hippocampus. Cell, 157(7), 1552–1564. https://doi.org/10.1016/j.cell.2014.03.067

Yu, H.-H., Chen, C.-H., Shi, L., Huang, Y., & Lee, T. (2009). Twin-spot MARCM to reveal the developmental origin and identity of neurons. Nature Neuroscience, 12(7), 947–953. https://doi.org/10.1038/nn.2345

Yu, Y.-C., Bultje, R. S., Wang, X., & Shi, S.-H. (2009). Specific synapses develop preferentially among sister excitatory neurons in the neocortex. Nature, 458(7237), 501–504. https://doi.org/10.1038/nature07722

