## Supplementary figures and images for "Sequential addition of neuronal stem cell temporal cohorts generates a feed-forward circuit in the Drosophila larval nerve cord"

### S1 Fig more twin spot data.pdf

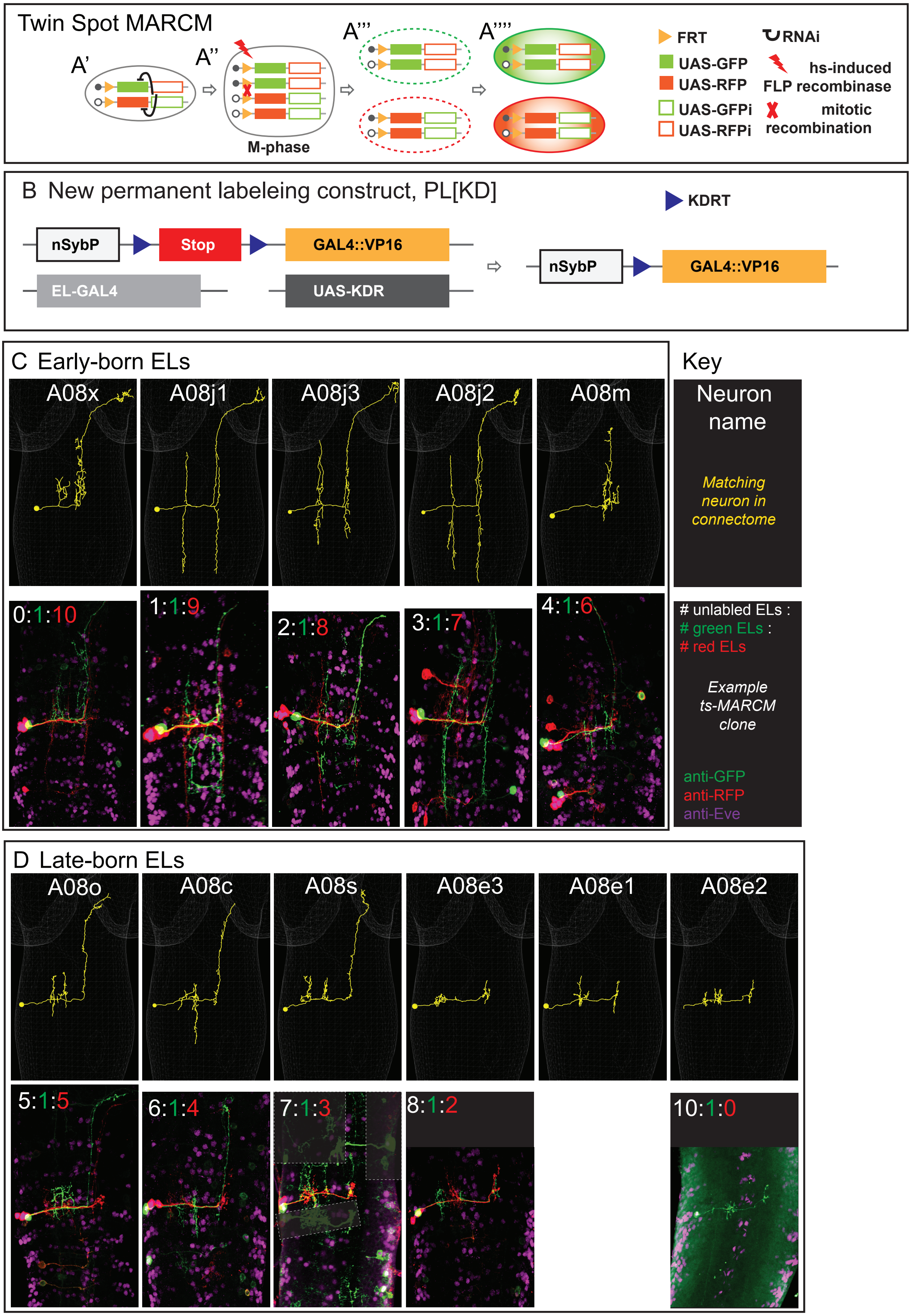


Supp Figure 1

### S2 Fig NB3-3 GAL4 and neurite lengths.pdf

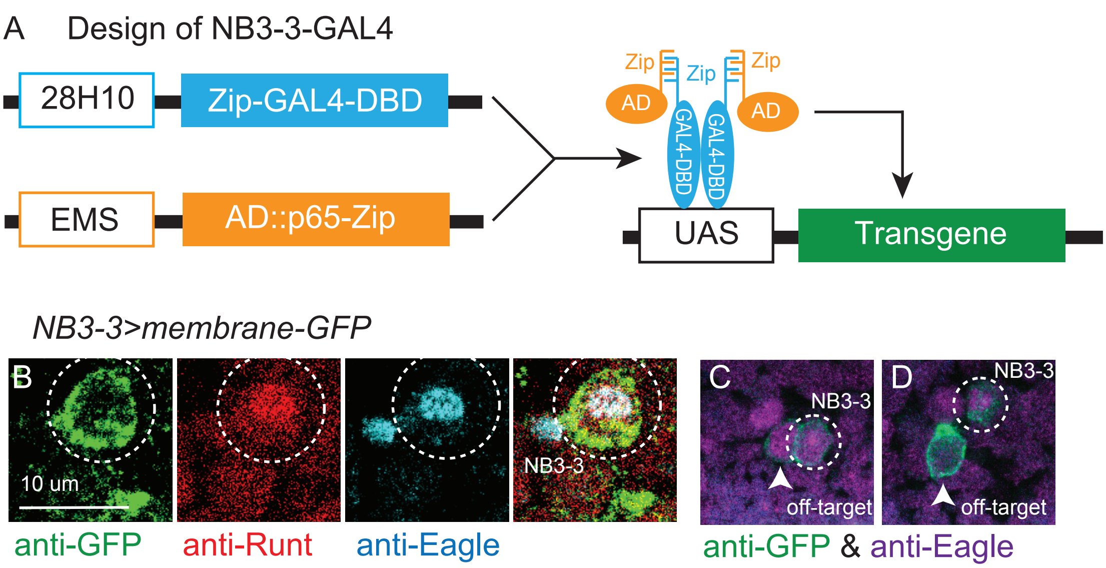


Supplemental Figure 2
